# Phosphoprotein dynamics of interacting tumor and T cells by HySic

**DOI:** 10.1101/2023.06.05.541905

**Authors:** Sofía Ibáñez-Molero, Jinne Pruijs, Alisha Atmopawiro, Fujia Wang, Maarten Altelaar, Daniel S. Peeper, Kelly E. Stecker

## Abstract

Functional interactions between cytotoxic T cells and tumor cells are central to anti-cancer immunity. Some of the proteins involved, particularly immune checkpoints expressed by T cells, serve as promising clinical targets in immunotherapy. However, our understanding of the complexity and dynamics of the interactions between tumor cells and T cells is only rudimentary. Here we present HySic (for <u>Hy</u>brid quantification of <u>S</u>ILAC (Stable Isotope Labelling by Amino acids in Cell culture)-labeled interacting <u>c</u>ells) as an innovative method to quantify protein and phosphorylation dynamics between and within physically interacting (heterotypic) cells. We show that co-cultured HLA/antigen-matched tumor and T cells engage in physical and stable interactions, allowing for in-depth HySic analysis. This method does not require physical separation of the two cell types for subsequent MS proteome and phosphoproteome measurement using label free quantification (LFQ). We demonstrate that HySic can be used to unravel proteins contributing to functional T cell:tumor cell interactions. We validated HySic with established interactions, including those mediating IFNγ signaling. Using HySic we identified the RHO/RAC/PAK1 signaling pathway to be activated upon interaction of T cells and tumor cells. Pharmacologic inhibition of PAK1 sensitized tumor cells to T cell killing. Thus, HySic is an innovative and simple method to study short-term protein signaling dynamics in physically interacting cells, which can be easily extended to other biological systems.

## Introduction

Immunotherapies, such as Immune Checkpoint Blockade therapy (ICB), have revolutionized cancer treatment. They act by targeting immune-suppressive protein interactions in the tumor microenvironment (TME). However, the majority of patients still fail to achieve a durable clinical benefit, due to therapy resistance, whether upfront or on therapy (Hellmann et al., 2019; Kalbasi & Ribas, 2020; Sharma et al., 2017). Cytotoxic T cells recognize tumor cells by TCR-antigen-MHCI interactions and subsequently trigger apoptosis. This inter-cellular communication serves as a critical event in the process of tumor eradication. Functional interrogation of T cell:tumor interactions using genetic approaches such as whole genome CRISPR-Cas9 screens (Kearney et al., 2018; D. W. Vredevoogd et al., 2021) has proven a successful strategy to identify genes vital to a tumor’s defense against T cell attack. For example, we previously showed that perturbation of specific components of pro-survival TNF signaling, like TRAF2 and RNF31, strongly sensitizes to killing by CD8 T and NK cells, *in vitro* and *in vivo* (Vredevoogd et al., 2019; Zhang et al., 2022). Together, these functional studies revealed several tumor resistance mechanisms, including lack antigen presentation and deficiencies in autophagy, IFNy and TNFa pathways (Kearney et al., 2018; Vredevoogd et al., 2019). However, such genetic approaches cannot expose resistance mechanisms that rely on dynamic protein networks and signal transduction events that are driven by post-translation modifications (PTMs), such as phosphorylation. This makes our understanding of the functional interactions between tumor and T cells rather gene-centric and rudimentary.

Phosphoproteomics techniques have allowed the identification of drug resistance mechanism in tumors (Boulos et al., 2020) and mechanisms of T cell key signaling molecules like TCR and PD-1 (Ruperez et al., 2012; Tocheva et al., 2020). The phosphoproteome has been shown to be a primary regulator of cell signaling during cell-cell interactions (Ardito et al., 2017) and in the last decade, accuracy in quantitative techniques has remarkably increased (Riley & Coon, 2016; Urban, 2022). However, standard proteomic and phosphoproteomic approaches quantify digested proteins from homogenized lysates, in which cellular context is lost during sample preparation. These methods, therefore, cannot distinguish the cellular origin of protein content, and do not allow for cell-specific information to be extracted from a mixed cell system such as T cells engaged with tumor cells. The need to investigate communication between cytotoxic T cells and tumor cells, two distinct cell types that physically interact, thus presents a technical challenge for (phospho)proteomic studies: preservation of cell-type-specific information, which is what we address here.

One approach to solve the challenge of deciphering mixed proteomes derived from interacting cells is the use of stable isotope labeling by amino acids in cell culture (SILAC) (Ibarrola et al., 2003; Ong et al., 2002). The proteome of different cells can be uniquely labeled by metabolic incorporation of essential amino acids that contain stable isotopes, thus providing a unique mass signature for each cell type. Traditional SILAC applications use labeling strategies to distinguish different experimental conditions by mixing labeled proteomes after cell lysis. The relative MS intensity of the labeled peptides are then used to quantify protein and phosphopeptide changes. This standard method utilizes SILAC for sample multiplexing and quantification during MS analysis. In addition, a handful of studies have applied SILAC labeling to study mixed cell types interacting in co-culture (Griffith et al., 2022; R. Liu et al., 2019). However, those studies either did not examine both interacting cell types, or they required physical separation by FAC-sorting after interaction, which can induce undesired changes in the phosphoproteome. Moreover, they utilize SILAC in the traditional way, as a ratio for quantification.

To overcome these technical limitations, here we developed an innovative strategy in which SILAC labels are used as barcodes to distinguish the two interacting cell types in co-cultures. A concomitant second method of label-free quantitation (LFQ) is applied to measure protein and phosphorylation dynamics for each SILAC signature separately. This hybrid-quantitative MS approach using both SILAC and LFQ allows for direct monitoring of both protein and phosphorylation dynamics over many different conditions, because we are not limited to a fixed set of ratios defined by SILAC. We termed this method <u>Hy</u>brid quantification of <u>S</u>ILAC (Stable Isotope Labelling by Amino acids in Cell culture) -labeled interacting <u>c</u>ells (HySic) to probe short-term proteome and phosphoproteome dynamics. To illustrate the utility of this method, we made use of a T cell:tumor cell co-culture system that we successfully used previously in genetic screens to identify genes critically involved in tumor sensitivity to T cells. Leveraging the standard SILAC labels, we deciphered three types of information in each experimental condition: protein degradation and phosphorylation occurring within T cells, within tumor cells, and the induction and phosphorylation of newly translated proteins upon co-culturing. After validating HySic findings from each of these categories using independent techniques, we explored T cell:tumor responses to find actionable targets. We further validated one target involved in both, T cell and tumor cell, phosphorylation responses. Altogether, we present the robustness and applicability of HySic as an easy platform to unravel protein dynamics and phosphorylation in functional (heterotypic) cell-cell interactions.

## Results

### Developing HySic

To study the signaling dynamics of T cell:tumor interactions, we made use of a system we established previously for melanoma and non-small cell lung cancer (NSCLC) (Ibañez-Molero et al., 2022). In this model, we genetically engineered tumor cells to express the major histocompatibility complex class I (MHC I, specifically HLA-A*02:01) protein capable of presenting the tumor antigen MART-1, as well as the MART-1 epitope itself. On the other hand, CD8^+^ T cells were isolated from healthy donors and subsequently retrovirally transduced with a MART-1-specific T cell receptor (TCR) (**Fig. 1A**). Consistent with our previous results, we observed that upon co-culturing, T cells were able to recognize the tumor cells. We validated that upon tumor cell encounter, T cells became activated, as judged by the production of cytotoxic molecules interferon γ (IFNγ) and tumor necrosis factor (TNF) (**Fig. 1B**). This ultimately led to tumor cell cycle arrest and/or cell death (**Fig. 1C).** These results illustrate the specificity and utility of this T cell:tumor co-culture model, providing the foundation to develop a method to analyze the functional interactions in more depth.

**FIG. 1.**
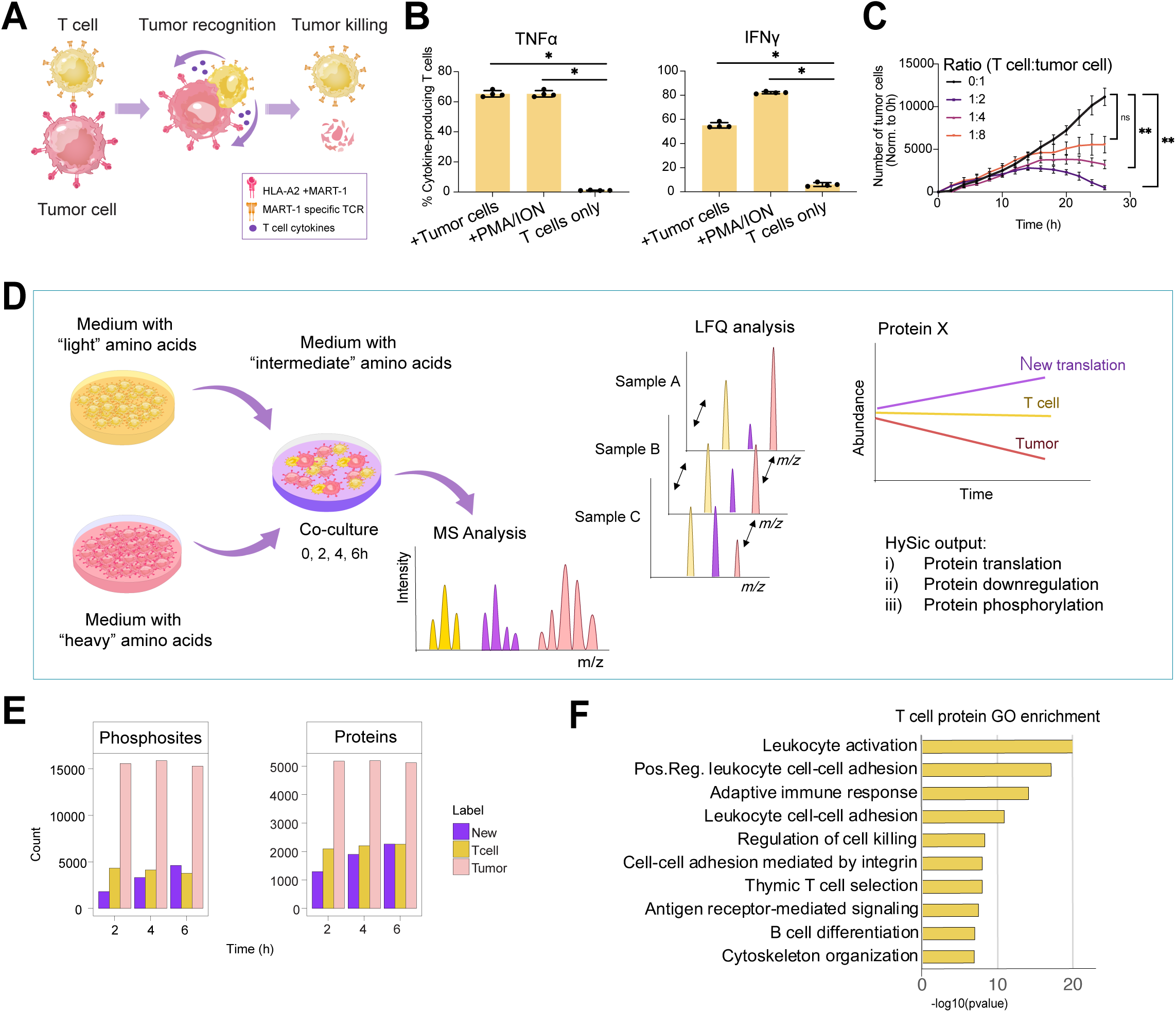
HySic: hybrid quantification of SILAC-labeled interacting cells. A) Schematic representation of T cell:tumor cell co-cultures. Tumor cells were transduced to express HLA-A2 and MART-1 antigen. T cells were isolated from PBMCs from healthy donors and subsequently transduced with virus encoding a MART-1-specific TCR. Matched interactions lead to tumor cell recognition and killing by the T cells. B) T cell activation after 4.5h co-culture as measured by TNFa and IFNy production by flow cytometry. Cells were co-cultured at 1:1 (T cell:tumor cell) ratio. As a positive control, T cells alone were cultured with PMA (50 ug/mL) and Ionomycin (1 mg/mL). As a negative control, T cells were not stimulated. Y axis represents % cytokine-producing T cells. Each dot represents a technical replicate. Error bars represent SD. Mann-Whitney test was used for analysis. C) Cytotoxicity assay of tumor cells (EBC-1) and T cells co-cultured at different T cell ratios as indicated. Viability of tumor cells (expressing mPlum) was measured by Incucyte tumor cell count (red fluorescence counts) every 2 hours. Data was normalized to 0h. Error bars represent SD of 3 technical replicates. Statistical analysis was performed by Friedman test. **p-value<0.01, ****p-value<0.0001. D) Schematic representation of HySic. T cells and tumor cells are labelled separately with “light” ^0^K^0^R or “heavy” ^8^K^10^R amino acids, respectively. Following complete label incorporation, the two cell types were co-cultured in medium with “intermediate” ^4^K^6^R amino acid isotopes. After 2, 4, and 6h, cells were lysed and analyzed by MS. Samples were subsequently quantified by LFQ by comparing extracted ion chromatograms for each SILAC label independently. Protein information was extracted from tumor cells, T cells or annotated as newly synthesized. E) Total count of quantified proteins and phosphosites by HySic with T cell, tumor cell or newly synthesized label, at the indicated time points. F) Top 10 enriched pathways of proteins enriched in the T cell labeled group. Proteins were selected for enrichment analysis if their expression in T cell label at T0 was > 2-fold higher than tumor label across time points. Pathways are listed from smallest to largest *p*-value. Enrichment of GO biological processes was performed using Metascape.

First, we tested whether this co-culture system allows for the measurement of protein and phosphorylation changes upon T cell:tumor engagement. After 16h, the T cell-containing supernatant was removed, followed by trypsinization of the fraction of tumor cells that were still conjugated to the T cells. This physical separation was performed in a panel of 15 human NSCLC cell lines, in which we observed that the purity of each cell fraction following separation was on average low: only 44% of the isolated T cell population was pure and did not contain tumor cells, and only 71 % of the isolated tumor cells were free from T cell contamination (**Fig. S. 1A**). This result indicates that there is a high percentage of contamination of the other cell type (whether tumor cell or T cell) in the separated fractions. This mixed cell content would complicate the interpretation of the proteomic and phosphoproteomic data by mass spectrometry (MS) analysis. To circumvent this problem, we developed a SILAC-based labelling method for analyzing functional interactions in the T cell:tumor cell co-cultures, as this could identify proteins of different cell origin from two cell mixed proteomes based on their unique mass signatures.

We labelled T cells and tumor cells separately with “light” (^0^K^0^R) and “heavy” (^8^K^10^R) amino acids, respectively. After >93% label incorporation was confirmed for four tumor cell lines (**Fig. S. 1B**), the two cell types were co-cultured for up to 6 hours in medium containing “intermediate” (^4^K^6^R) amino acids isotopes. Cells from complete co-cultures were then pelleted, lysed and analyzed by LC-MS for proteome and phosphoproteome analyses. The incorporation of three different amino acid isotope labels allows for the potential identification of proteins from distinct origins: tumor cells (^8^K^10^R), T cells (^0^K^0^R) and newly synthesized proteins from either cell type (^4^K^6^R). We will refer to these isotope labels as tumor-label, T cell-label and newly synthesized-label, respectively. In this way, we exploited SILAC labels as barcodes to distinguish the two cell types. We next used a second MS quantification method, label-free quantification (LFQ), to measure the change in protein and phosphosite abundance for each SILAC label independently across samples (**Fig. 1D**). Together, this constitutes a <u>Hy</u>brid quantification approach to measure <u>S</u>ILAC-labeled **i**nteracting <u>c</u>ells (HySic). We applied HySic to analyze four different NSCLC cell lines in cultured with T cells, as a function of time (0, 2, 4 and 6 hours) and at a 1:1 T cell:tumor cell ratio. We conducted these experiments in two independent batches. To control for background translation, we also incubated tumor cells and T cells independently in newly synthesized-label media for 4 hours.

We first analyzed the total number of unique proteins and phosphosites quantified by HySic (**Fig. 1E**). In tumor cell fractions, we measured more proteins and phosphosites than in the T cell or newly synthesized fractions. This was likely due to a higher protein content in tumor cells owing to their bigger size (average 22,9 µm diameter compared to 6,9 µm of T cells). We also observed an increase in proteins and phosphoproteins in the newly synthesized channel over time, which was expected because of new protein translation. To confirm the specificity of our SILAC labels, we analyzed proteins that had higher abundance in the T cell-label and observed an enrichment in leukocyte activation, leukocyte cell-cell adhesion and adaptive immune response, amongst other T cell-related pathways (**Fig. 1F**). This result confirms that proteins detected in the T cell-label are of T cell origin, demonstrating the specificity and efficiency of SILAC-barcoding of the HySic differential labelling method.

### HySic identifies interaction-induced newly synthesized proteins

We next used HySic to analyze the newly synthesized proteins, identified by the newly synthesized-label. To control for baseline translation, we compared the newly synthesized proteins in co-culture to individual incubation of tumor cells in newly synthesized-label media for 4 hours. We searched for proteins that were exclusively upregulated in co-culture and extracted the newly synthesized proteins that were shared across cell lines for pathway enrichment analysis. Proteins were enriched in Type II interferon signaling (IFNγ) and cell death (**Fig. 2A**), which is indicative of a TCR-antigen-MHCI interaction triggering T cell secretion of IFNγ causing subsequent tumor cell death (Jorgovanovic et al., 2020). These findings validate our ability to quantify expected biological responses using our HySic approach.

**FIG. 2.**
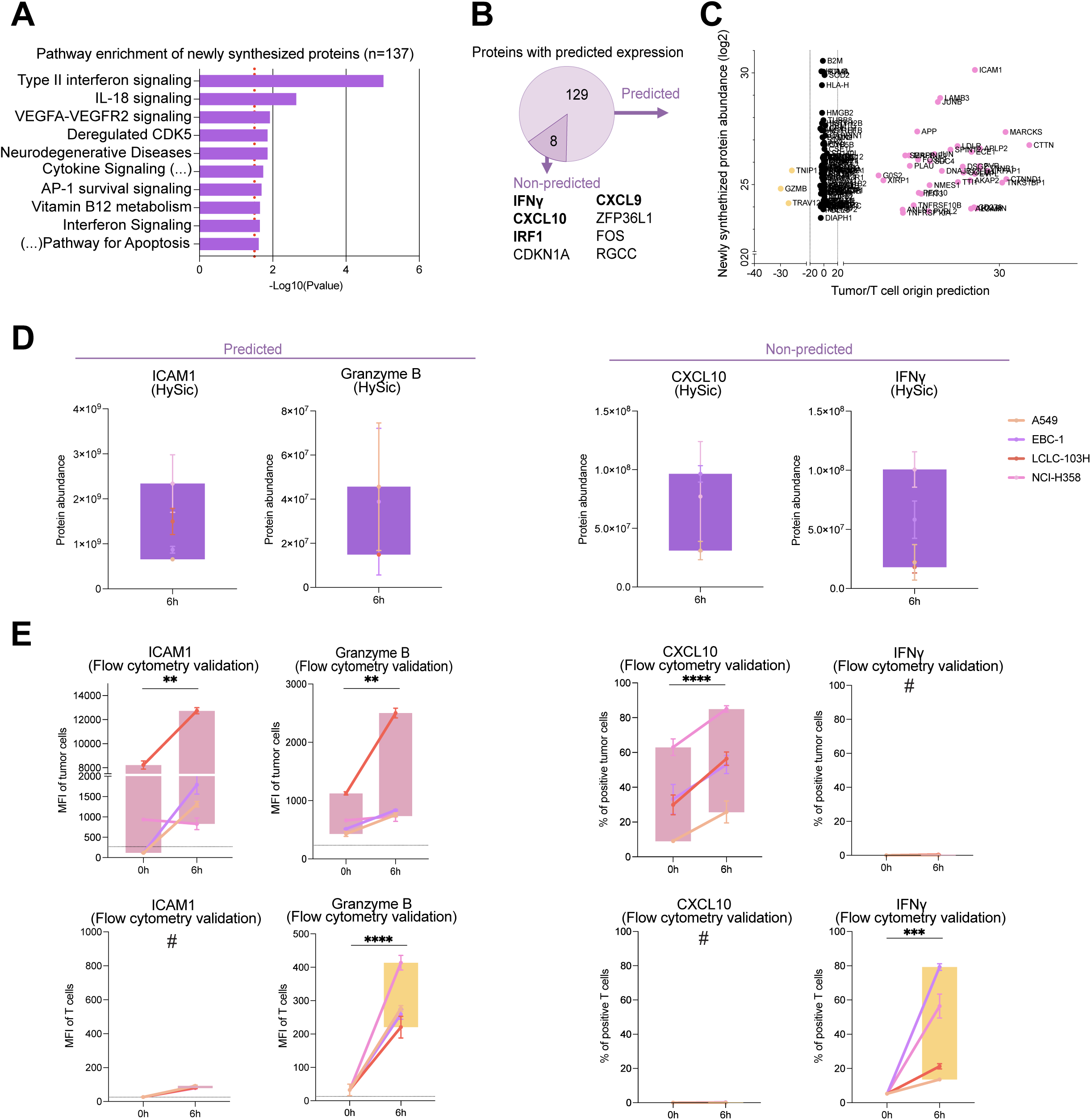
HySic identifies interaction-induced newly synthesized proteins. A) Top 10 enriched pathways in proteins detected in the channel for newly synthesized proteins. Enrichment was performed using *Gprofiler* with Reactome and WikiPathways databases. Pathways are listed from smallest to largest *p*-value. B) Protein expression prediction diagram. Of the 137 detected newly synthesized proteins across cell lines, 129 proteins were predicted to have T cell or tumor origin, whereas 8 proteins could not be predicted due to lack of expression at baseline in both cell types. C) Protein prediction scores of 137 newly synthesized proteins. Tumor/T cell prediction score in X axis was calculated as the difference of log2(Tumor/T cell) expression of the indicated protein at baseline. A positive score indicate Tumor predicted expression, whereas a negative score indicate T cell predicted expression. Y axis indicates log2(protein abundance) of the protein in the newly synthesized channel. D) Protein abundance detected by HySic of the indicated proteins. Each cell line is represented by a colored dot with SD bars. Boxplots indicate deviation of four cell lines combined. E) Median fluorescence intensity (MFI) or percentage of positive cells measured by flow cytometry of the indicated proteins in either tumor cells (pink boxplots) or T cells (yellow boxplots). Each cell line is represented by a colored dot with SD bars. Boxplots indicate deviation of four cell lines combined. **p-value<0.01, ****p-value<0.0001. 2-way ANOVA test used for statistical analysis. # indicates below detection levels.

In co-cultures, the cell origin of newly synthesized proteins is unknown, as tumor cell- and T cell-translated proteins share the newly synthesized-label. To tackle this limitation, we established a prediction method based on the baseline protein expression in tumor and T cells. A tumor/T cell prediction score was determined by dividing the baseline expression of a protein in each given cell type. A positive score indicates tumor origin, and a negative one T cell origin. Of the 137 newly synthesized proteins, 129 were predicted, while for 8 we were unable to make an assignment (non-predicted) due to lack of expression at baseline in either cell type (**Fig. 2B**). Within the non-predicted translated proteins, we identified known mediators of T cell:tumor cell interactions, including CXCL10, CXCL9, IFNγ and IRF1 (Benci et al., 2016). Among the predicted proteins, we observed more tumor cell origin assignment (**Fig. 2C**), likely due to the higher overall tumor cell protein content measured (**Fig. 1E**). The predicted newly translated proteins included those of known tumor origin like ICAM1 (Bui et al., 2020; Kotteas et al., 2014) and CDH1 (Abascal et al., 2016); and known T cell origin, GZMB and TRAV12 (Lieberman, 2003), corroborating our approach.

We next wanted to validate our HySic results for new protein translation using an independent technique. We selected non-predicted proteins (IFNγ and CXCL10) and predicted proteins in tumor (ICAM1) and T cells (GZMB) (**Fig. 2D**) and evaluated their protein expression in both cell types using flow cytometry. After co-culturing, IFNγ and GZMB expression increased in T cells, whereas CXCL10 and ICAM1 expression increased in tumor cells, as anticipated (**Fig. 2E**). Interestingly, GZMB protein expression was also increased in tumor cells upon interaction with T cells, in agreement with a recent study (Jiang et al., 2020). These results were reproduced with an independent T cell donor in the four cell lines (**Fig. S. 2A**). As a negative control, we evaluated proteins that did not show induced translation upon coculture. We evaluated PD-L1, which was not increased upon T cell co-cultured in three of the four tumor cell lines. We confirmed the absence of newly translated PD-L1 in those cell lines (**Fig. S. 2B**), validating the quantitative nature of our assay. Together, these data show that HySic in T cell:tumor cell co-cultures can be used to identify and quantify newly synthesized proteins upon cell:cell interactions.

### T cell:tumor cell interactions promote turnover of CD8A, Granulins and EGFR

Another type of protein information obtained by HySic is protein downregulation. By labelling the proteins derived from either tumor cells or T cells we can track their decrease during the co-culture, in comparison with individual cell populations. We evaluated the T cell and tumor cell protein expression in co-culture over time. We normalized the data to cells cultured individually to correct for generic protein turnover. Then, the dataset was filtered for proteins that were significantly downregulated, relative to timepoint zero, within at least one co-cultured cell line and showed downregulation across at least three of four cell lines (**Fig. S.3A and B)**. We performed pathway enrichment analysis to identify cellular processes affected by protein downregulation and observed that downregulated proteins in T cells were enriched in immune system pathways mainly, while in tumor cells they were enriched in a wider variety of processes (**Fig. S. 3C**). We next filtered the data for proteins decreasing across all six hours co-culture incubations and ranked the average protein downregulation (**Fig. 3A and B**). From the top 20 proteins with the largest average downregulation, we selected proteins for validation based on antibody availability: CD8 in T cells, and GRN and EGFR in tumor cells. CD8 was previously shown to be downregulated upon TCR-antigen-MHCI interaction (Xiao et al., 2007). To mimic the procedure of HySic, we pre-labelled CD8 and EGFR with a flow cytometry antibody and co-incubated T and tumor cells for 6 hours, after which, the remaining labelled protein was measured. Indeed, both CD8 and EGFR protein levels were decreased after co-culture in four independent tumor cell lines, with the decrease in EGFR being of less magnitude than CD8 (**Fig. 3C)**, validating the observations made and illustrating the quantitative power of HySic. These results were reproduced with an independent T cell donor (**Fig. S. 3D**). Using an alternative approach, we evaluated GRN protein expression by western blotting before and after T cell:tumor co-culture. We observed a decrease in GRN expression upon T cell co-culture in three of four cell lines (**Fig. S. 3E-G**), which supports our observed downregulation of GRN in tumor cells upon T cell interaction.

**FIG. 3.**
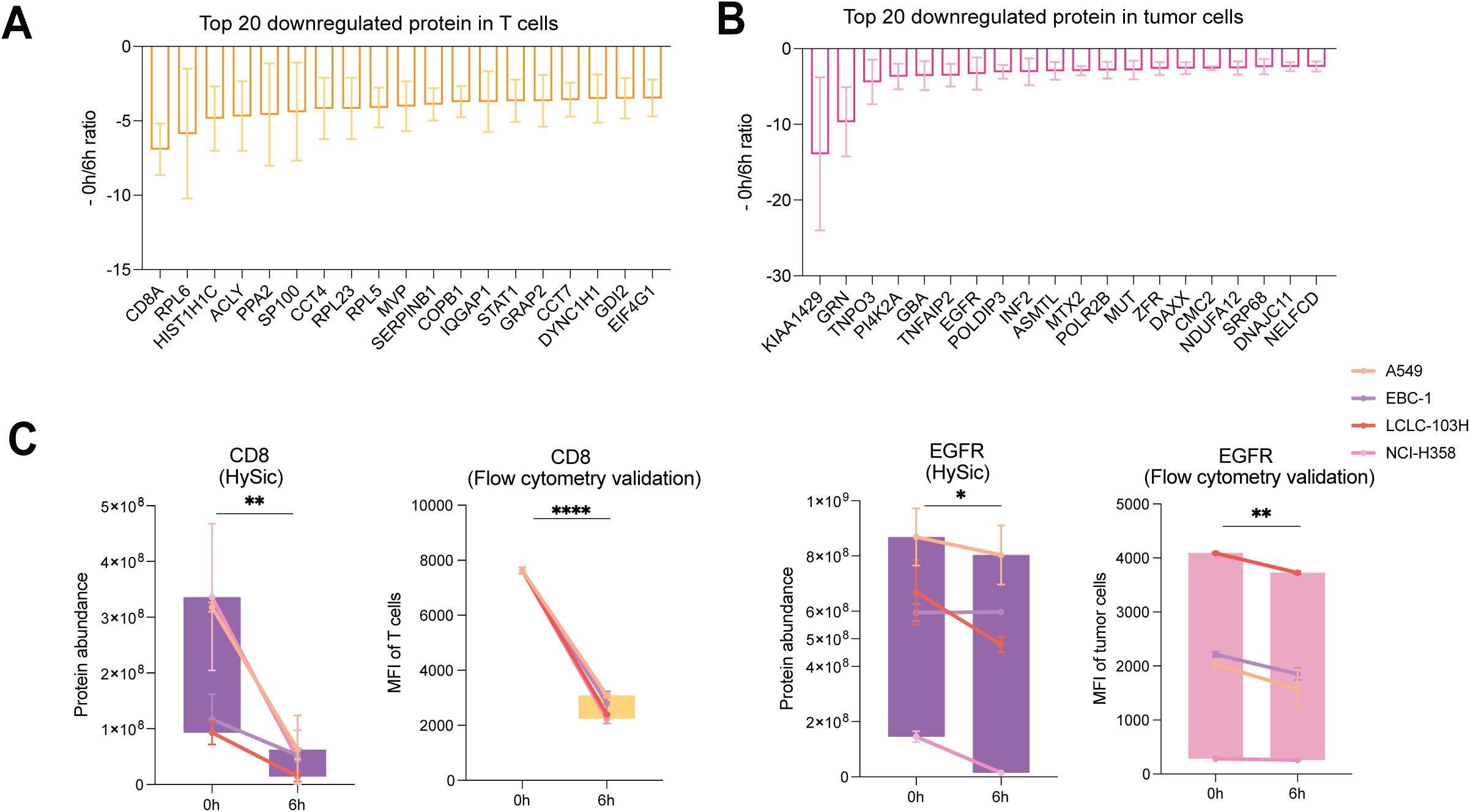
T cell:tumor cell interactions promote turnover of CD8A, Granulins and EGFR. A) Rank of top 20 downregulated proteins selected in T cell shown on X axis. Each dot represents a cell line, errors bars represent SEM. Y axis represent negative 0/6h protein expression ratio. B) Rank of top 20 downregulated proteins selected in tumor shown on X axis. Each dot represents a cell line, errors bars represent SEM. Y axis represent negative 0/6h protein expression ratio. C) Abundance of CD8 and EGFR proteins (purple) detected by HySic. Median fluorescence intensity (MFI) measured by flow cytometry of the indicated proteins in either T cells (yellow boxplots) or tumor cells (pink boxplots). Each cell line is represented by a colored dot with SD bars. Boxplot indicates deviation of four cell lines combined. *p-value<0.05, **p-value<0.01, ****p-value<0.0001. 2-way ANOVA test used for statistical analysis.

### HySic identifies induced RHO/RAC/PAK1 signaling upon T cell:tumor cell interactions

In addition to protein translation and degradation, the HySic approach allows for the measurement of phosphorylation dynamics between and within interacting cells. Protein phosphorylation is the most common post-translational modification and it heavily influences the function of proteins and protein networks (Ardito et al., 2017; Pawson & Scott, 2005). Therefore, we set out to establish which cellular processes are changed upon cell:cell interactions at the phosphorylation level within tumors, T cells, and in newly translated proteins. We first examined the newly synthesized phosphoproteins and found nearly 100 phosphosites upregulated > 2-fold above background translation at later timepoints (4h and 6h) in at least three of four co-cultured cell lines (**Fig. S. 4A**). These phosphorylated proteins showed high enrichment for factors associated with TNF signaling (**Fig. S. 4B)**, which can be explained by the important role of TNF in mediating immune cell cytotoxicity. This finding contrasts with the IFNG pathway that was mainly enriched at the level of protein translation (**Fig 2A**). Taken together, these data demonstrate the unique layers of information that can be revealed by the combination of proteome and phosphoproteome data.

We next performed a hierarchical clustering of the significantly regulated phosphorylation sites for tumor cells and T cells that were shared across at least three out of four of the co-cultured cell lines (**Fig. 4A**). The clusters containing increased phosphorylation were analyzed for pathway enrichment. The tumor cells and T cells revealed a shared pathway enriched for RHO/RAC signaling (**Fig. 4B and 4C**). RHO/RAC signaling includes multiple downstream cellular processes, many of which are linked to both cancer progression (Clayton & Ridley, 2020; Lin & Zheng, 2015) and T cell migration and activation (Saoudi et al., 2014). There are five major downstream RHO/RAC effector proteins: ROCK (1-2), PAK (1-6), mDia, WASP and WAVE (Møller et al., 2019). To understand which of these accounts for the downstream signaling in our model, we analyzed the phosphorylation status of the five candidates upon T cell:tumor cell co-culture in four tumor cell lines (from which phosphosites were detected in MS). The phosphorylation coverage of several candidate effector proteins was sparse and showed varied patterns of regulation. However, amongst all tested proteins, PAK1 activating sites (pS144, pS174, pT212, pS223) were consistently more phosphorylated upon T cell:tumor cell interactions (**Fig. 4D, E and Fig. S. 5A, B**). WAVE also showed activating phosphosites upregulated across different cell lines. Conversely, ROCK1 showed decreased phosphorylation of the activating site pS1341.

**FIG. 4.**
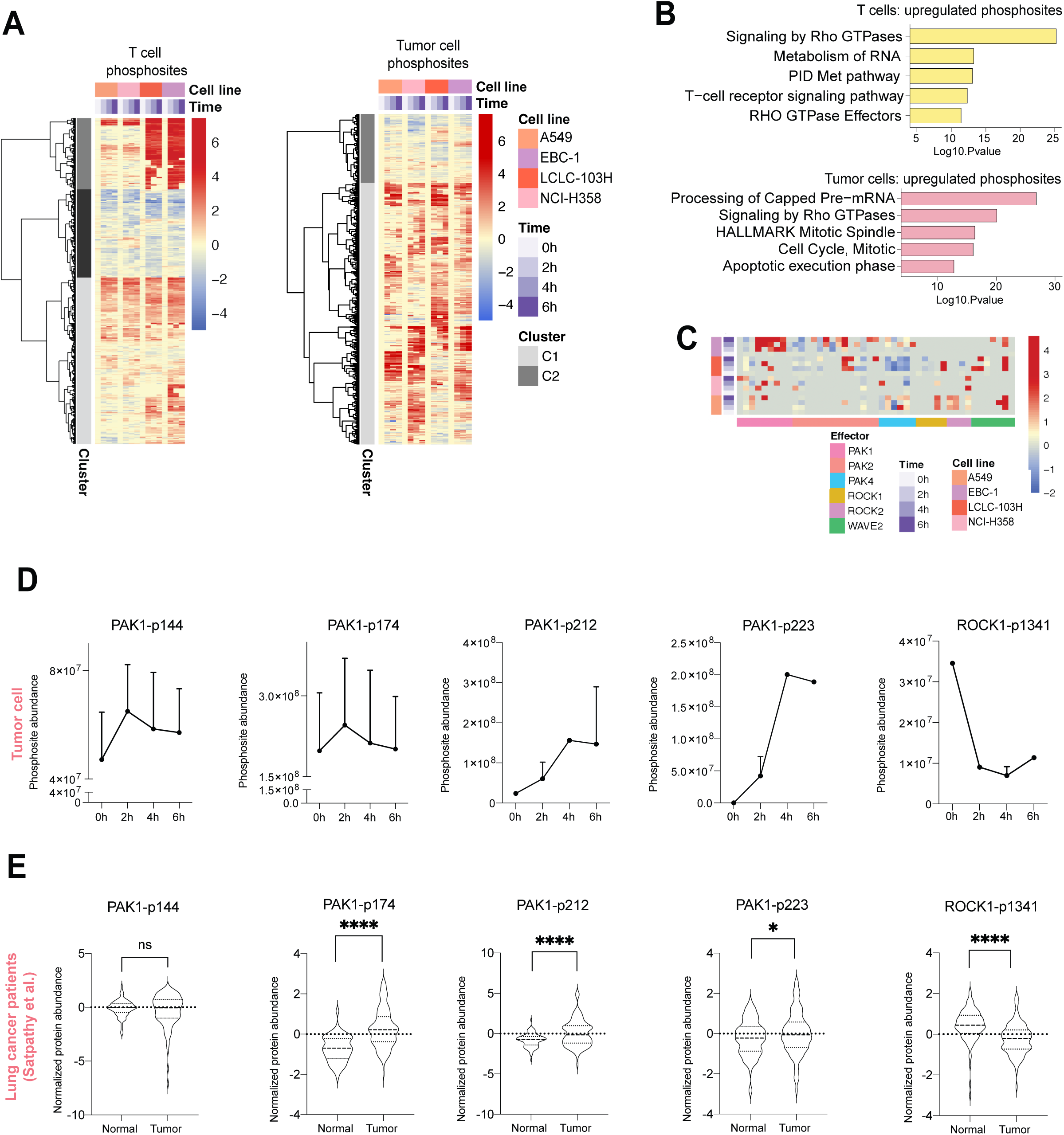
T cell:tumor cell interaction upregulates RHO/RAC signaling and PAK1 activation phosphosites, that are also increased in tumor vs normal tissue. A) Significantly regulated phosphosites (p<0.05) shared across at least 3/4 co-cultured cell lines for T cells (left) and tumor cells (right). Phosphosite expression values represent T0 normalized data. Missing values are replaced with zero for heatmap clustering. B) Pathway enrichment (by Metascape) for significantly upregulated phosphorylated proteins from phosphosite heatmap clusters in T cells (C1&C3) and tumor cells (C2). Top five enrichment terms are displayed. C) Heatmap clustering of Rho/Rac effector protein phosphorylation sites. Missing values are replaced with zero for heatmap clustering. D) Phosphosite abundance detected by HySic of the indicated protein sites in tumor cells. Each dot represents average of four cell lines and error bars represent SEM. E) Normalized phosphosite abundance from Satpathy *et al*. of the indicated protein sites in normal versus tumor tissue. Violin plots represent average of individual patient expressions.

To start exploring whether the phosphorylation of activating sites in PAK1 and ROCK1 could have clinical relevance, we analyzed the same phosphosites in a lung cancer clinical study, which includes phosphoproteomics of tumor and healthy tissue (Satpathy et al., 2021). PAK1-activating phosphosites were more abundant in tumor compared to healthy tissue (**Fig. 4E**). This was reproduced in an independent lung cancer cohort (Gillette et al., 2020) (**Fig. S. 6A**). ROCK1-activating phosphorylation was, by contrast, lower in tumor vs healthy tissue, following the trend observed in T cell:tumor co-cultures. The remaining effector proteins were also evaluated, but no consistent trends were observed, or data was not available (**Fig. S. 6B**). The increase in activating phosphorylation in PAK1 upon T cell:tumor cell engagement led us to investigate PAK1 as a possible target in tumor immunotherapy.

### PAK1 inhibition enhances tumor cell sensitivity to T cells

PAK1 has been previously studied in cancer (Chow et al., 2022; Chung et al., 2020; H. Liu et al., 2021; Semenova & Chernoff, 2017; Wang et al., 2020), but its role in the context of cytotoxic T cell killing has not yet been examined. Therefore, we first wished to increase our understanding of the role of PAK1 in T cells, and second evaluate its action in T cell:tumor co-cultures. To inhibit PAK1 activity, we treated T cells with a selective inhibitor (NVS-PAK1) (**Fig. S. 7A**). We first evaluated the on-target effect of the inhibitor by phosphoproteomic analysis of T cells treated for one or five days with NVS-PAK1. We observed a decrease in the PAK1 auto-phosphorylation site pS144 (**Fig. S. 7B**). This phosphosite serves as a positive control for PAK1 inhibition as PAK1 must be functional to perform S144 autophosphorylation (Chong et al., 2001). We next analyzed protein expression changes in T cells upon NVS-PAK1 treatment. At the proteomic level, we observed an increased in the T cell-derived cytotoxic protein Granzyme B (**Fig. 5A**). We validated the increase in GZMB protein expression by flow cytometry upon treatment with NVS-PAK1, as well with a second PAK1 inhibitor, FRAX-597 (**Fig. 5B and S. 7C**). In addition, we evaluated the effect of PAK1 inhibition on T cells by analyzing T cell activation markers by flow cytometry. Activation markers CD69 and CD137 were increased upon NVS-PAK1 treatment in T cells (**Fig. S. 7D**). Thus, PAK1 inhibition on T cells increases the production of the cytotoxic molecule GZMB and increases the expression of cell surface activation markers.

**FIG. 5.**
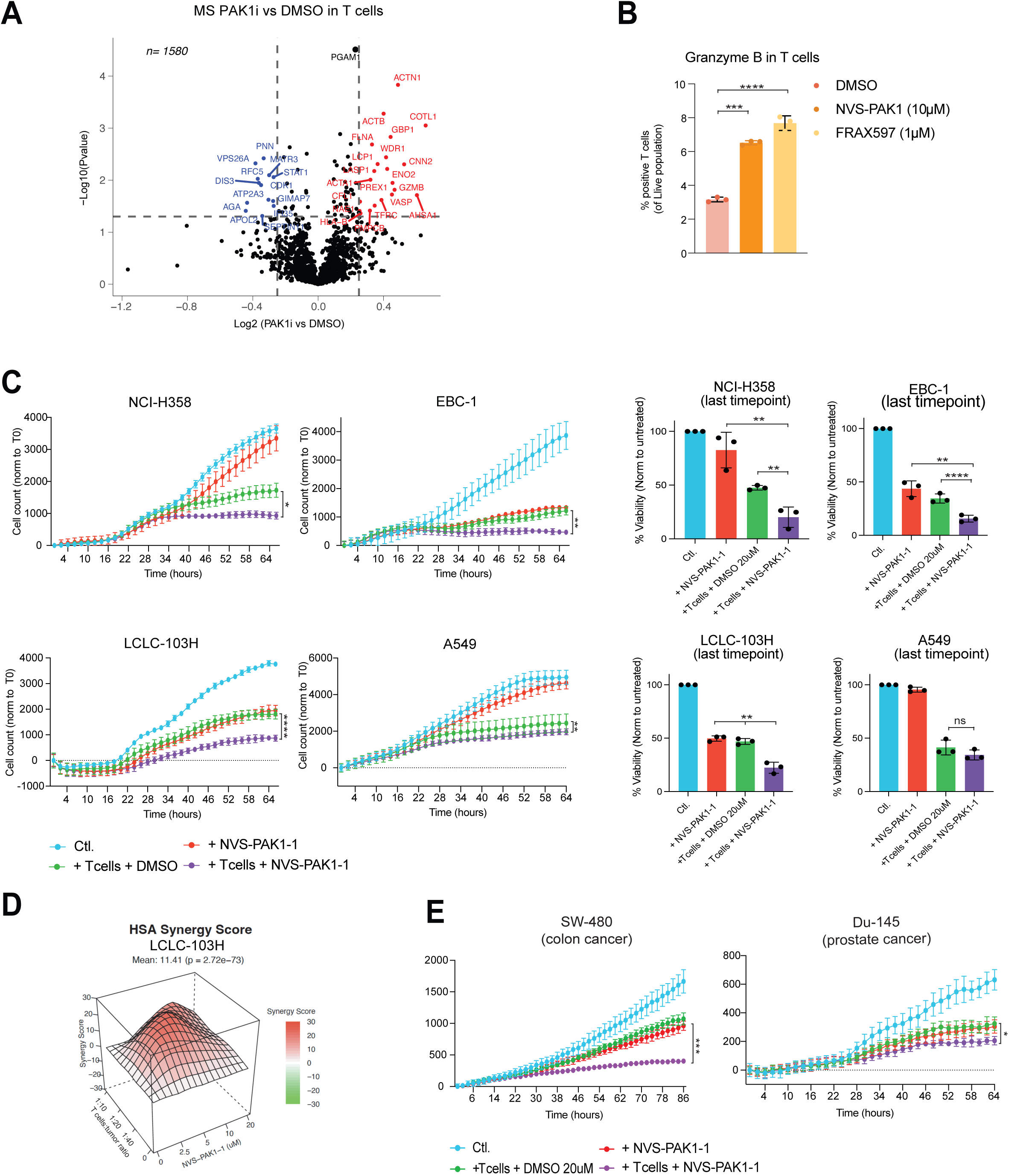
Inhibition of PAK1 increases T cell sensitivity. A) Proteome profiling of NVS-PAK1-treated (10 uM, 5d) vs DMSO-treated T cells. Colored dots are significantly increased (red) or decreased (blue) by NVS-PAK1 treatment (p< 0.05) with a log2 fold-change greater than 0.25. B) Percentage of Granzyme B-positive T cells (of live population) upon treatment with NVS-PAK1 (10uM), FRAX587 (1uM) or DMSO for 5 days. ***p-value<0.001, ****p-value<0.0001. 2-way ANOVA test used for statistical analysis. C) (Left) Cytotoxic assay of tumor and T cells co-cultured with or without NVS-PAK1 in Incucyte®. Inhibitor concentration and T cell:tumor ratio was optimized for each cell line; NCI-H358 (2.5uM and 1:20), EBC-1 (10uM and 1:10), LCLC-103H (5uM and 1:10) and A549 (2.5uM and 1:10). Viability of tumor cells (expressing mPlum) was measured by red fluorescence counts every 2 hours. Data was normalized to 0h. Error bars represent SD of 3 technical replicates. Statistical analysis was performed by Friedman test. *p-value<0.05, **p-value<0.01, ****p-value<0.0001. (Right) Endpoints of experiments shown left, normalized to untreated control. Each dot represents a biological replicate and error bars represent SD. Nested T test was used for statistical analysis. **p-value<0.01, ****p-value<0.0001. D) Synergy score (HSA) calculated with *Synergyfinder* on the indicated T cell:tumor cell ratios and drug concentrations for LCLC-103H cells. Score indicates the percentage of additive effect of using a combination of two treatments compared to the single agents. E) Cytotoxic assay of tumor and T cells co-cultured with or without NVS-PAK1 in Incucyte®. Inhibitor concentration and T cell:tumor ratio were optimized per cell line; SW480 (20uM, 1:10), DU-145 (10uM, 1:10). Data was normalized to 0h. Error bars represent SD of 3 technical replicates. Statistical analysis was performed by Friedman test. *p-value<0.05, ***p-value<0.001.

Lastly, we set out to determine whether PAK1 inhibition would have a beneficial effect on T cell-mediated tumor killing. T cells and tumor were co-cultured and treated with NVS-PAK1 at various T cell:tumor ratios in a time-course analysis, in four independent cell lines. NVS-PAK1 treatment significantly increased tumor sensitivity to T cell killing in three out of the four NSCLC cell lines tested (**Fig. 5C**). The effect of this drug was additive to T cell killing in two NSCLC cell lines (score <10 in EBC-1 and NCI-H358), and synergistic in the third cell line (score >10 in LCLC-103H) as evaluated by has synergy score. In the fourth cell line there was no additive nor synergistic effect observed (**Fig. 5D 7E**). To extend these findings to other cancer types, we repeated the experiment in breast, colon and prostate cancer cell lines. NVS-PAK1 again significantly increased T cell killing sensitivity in colon and prostate cancer cell lines, but not in the breast cancer cell line (**Fig. 5E and S. 7F**). Together, these data indicate that PAK1 inhibition can enhance the sensitivity of tumor cells to T cell killing in multiple cancer types, including NSCLC, colon and prostate cancer.

## Discussion

Deciphering the complex signaling networks that mediate a tumor cell’s defense against T cell attack requires comprehensive analyses, particularly at the protein level. However, such (phospho)proteomic studies have been limited owing to the technical challenge of preserving cell-type-specific information in a mixed-cell system. Here, we have developed a method to determine short-term protein translation and phosphorylation dynamics in interacting tumor and T cells: HySic. Although previous studies performed proteomics of interacting cells, mainly by the use of SILAC (Griffith et al., 2022; R. Liu et al., 2019; Ramello et al., 2019), they either made use of physical cell:cell separations, or did not study the individual cell types involved in the interaction. Therefore, we developed a workflow allowing for SILAC-labeled cells to interact without the need of a physical separation step, leaving the (phospho)proteome landscape unaltered. We simultaneously analyzed proteins and phosphosites derived from both tumor and T cells, as well as the proteins that were newly translated at the time of their interactions. SILAC was used to identify the origin of a given protein/phosphosite, followed by label free quantification of the individual SILAC labeled peptides. This combined data analysis allowed us to track proteins and phosphosites dynamics over different cell types, time and treatment conditions. We validated data for protein translation and degradation obtained by HySic, while our integrative phosphoproteomics analysis identified PAK1 as a potential target for tumor sensitization to T cell killing.

As one aspect of our method, we studied protein degradation upon T cell:tumor interaction. We identified proteins that were downregulated upon cell interaction, and we selected several for validation. As an expected event, we observed the turnover of CD8 in T cells upon T cell:tumor cell interaction (Xiao et al., 2007). On the tumor cell side, we observed the downregulation of EGFR. EGFR inhibition has been shown to correlate with increased immune infiltration in cancer patients (Li et al., 2019; Soucheray et al., 2015). Following this and our proteome observations, the inhibition of EGFR may be involved in the tumor response to T cell attack, but this needs further study. To confirm the downregulation of EGFR, we analyzed EGFR phosphosite expression and observed the increase of p1039, p1041, and p1042, which are known to be involved in the receptor internalization (**Fig. S. 8A**). We also observed downregulation of granulin, a mitogen involved in tumor proliferation and invasion (Arechavaleta-Velasco et al., 2017; Zanocco-Marani et al., 1999). Recently, it has been shown that the granulin precursor (Pro-granulin) can induce PD-L1 and that blocking Pro-granulins can sensitize tumors to T cell killing in an immunocompetent mouse model (Cheung et al., 2022; Fang et al., 2021). The mechanisms behind these and other downregulated proteins in interacting tumor and T cells remain to be further studied.

In addition to protein translation and turnover, HySic enables the quantification of cell-specific phosphoproteomics data. Following co-culture, tumor and T cells are snap frozen together, without additional perturbation caused by cell-separation methods, which can profoundly influence phosphorylation dynamics. This allows capturing more transient phosphosites that could change within seconds otherwise (Blazek et al., 2015). By analyzing the most abundant changes in the phosphoproteome of both tumor and T cells, we found a shared pathway being altered during cell-cell engagement: RHO/RAC signaling. This pathway is broadly involved in cell-cell communication via modulation of the cell cytoskeleton, amongst other functions (Burridge & Wennerberg, 2004) One of its effector proteins, PAK1, has been thoroughly studied in tumor cells. We observed that activating phosphosites of PAK1 were increased in tumor versus adjacent normal tissue in cancer patients. However, considerably less is known about the role of PAK1 in T cells, or in the context of T cell:tumor interactions. By pharmacologically inhibiting PAK1, we uncovered new properties associated with this effector protein in T cells: induction of Granzyme B and increased expression of activation proteins. Furthermore, we show that PAK1 inhibition increases tumor sensitivity to T cell killing.

PAK1 is already being considered as an attractive oncological target, due to its contribution to several oncogenic signaling pathways in cancer cells (Semenova & Chernoff, 2017). Most described PAK1 inhibitors also target other PAK family members, which can lead to toxicity. Recently, a more selective inhibitor (NVS-PAK1, used in this study) was developed, but its efficacy in an *in vivo* Schwannoma mouse model was limited (Hawley et al., 2021). Future development of an efficient and selective PAK1 inhibitor could help to translate our findings in pre-clinical models and perhaps the clinic. Following the functional validation of PAK1, other phosphoprotein pathways described may also be explored to boost mechanisms of tumor sensitivity to T cells.

One limitation of HySic concerns the analysis of newly synthesized proteins. In our study, both tumor and T cells were co-cultured in ^4^K^6^R labelled medium, which does not allow us to distinguish the cell origin of newly synthesized proteins. To find a solution for this, we implemented a prediction method based on the baseline expression of proteins on each cell type. We were able to assign a prediction score to the majority of proteins and validate several of them in both tumor and T cells. However, for any newly synthesized protein of interest, additional validation is required to confirm the cell of origin.

Additionally, co-culture incubation times using the HySic approach must be kept short (<8 hours) to avoid extensive proteome mixing due to protein turnover. To avoid proteome mixing in SILAC co-cultures, cell lines can be genetically engineered to uniquely metabolize and incorporate amino acid precursors (Gauthier et al., 2013). This approach has been successful for delineating translation in mixed cell cultures. However, it requires an extensive experimental setup involving exogenous expression of metabolic enzymes, which can be difficult to implement, especially in primary cells like T cells. Therefore, we view HySic as a simple and easy alternative for studies prioritizing short-term signaling dynamics between interacting cells.

In conclusion, we have established HySic as an MS method to identify protein and phosphoprotein changes in SILAC-labeled interacting cells. After validation of the method, we have explored the possibilities of the technique. HySic allowed identifying newly translated and downregulated proteins, as well as mapping phosphoprotein dynamics. The latter option enables us to identify and explore PAK1 as a target to increase tumor sensitivity to T cells. Beyond PAK1, also other pathways and actionable targets have emerged from our dataset, which merits further investigation. HySic is a relatively easy method that can retrieve extensive functional protein data in different co-cultured cell types.

## Material and Methods

### Cell culture

Tumor cell lines were obtained from the Peeper laboratory stock. They were cultured in RPMI (Thermofisher, 21875034) supplemented with 10% fetal bovine serum (Sigma, 3101120) and 100U/ml Penicillin-Streptomycin (Invitrogen, 15140-122). They were transduced with the HLA-A*02:01-MART1-mPlum lentiviral plasmid as described previously (Ibáñez et al. manuscript under revision). They were tested for mycoplasma by PCR monthly.

### SILAC labelling

Tumor cell lines were cultured in “Heavy” ^8^K^10^R medium consisting of RPMI 1640 Medium for SILAC (88365, Thermofisher) with 10% dialyzed fetal bovine serum (Fisher Scientific, 15605639), 100U/ml Penicillin-Streptomycin (Invitrogen, 15140-122), 0.04 mg/L proline (Sigma Aldrich, P5607), 100ug/ml L-arginine:HCL 13C6, 99%; 15N4, 99% (Cambridge isotope laboratories, CNLM-539-H-0.25) and 40mg/ml L-lysine:2HCL 3,3,4,4,5,5,6,6-D8, 98% (Cambridge isotope laboratories, CNLM-539-H-0.25). Cells were grown in this medium for at least 5 doublings before testing isotope incorporation by MS.

### T cell:tumor co-culture with SILAC

CD8 T cells were isolated and transduced as described previously (David W. Vredevoogd et al., 2019). Briefly, CD8 T cells were isolated with Dynabeads from PBMCs from healthy donors, activated with aCD3 and aCD28 antibodies (eBioscience, 5 µg per well) for 48h and transduced with a MART-1 specific TCR by spinfection. Stable-isotope labelled tumor cells were counted and seeded at 3×10^6^ cells per 10 cm petri dish. The next day, 3×10^6^ MART-1-transduced T cells were added and incubated for the indicated hours (2, 4 or 6) in “Intermediate” ^4^K^6^R medium consisting of RPMI 1640 Medium for SILAC (88365, Thermofisher) with 10% dialyzed fetal bovine serum (15605639, Fisher Scientific), 100U/ml Penicillin-Streptomycin (Invitrogen, 15140-122), 0.04 mg/L proline (Sigma Aldrich, P5607), 100ug/ml L-arginine:HCL 13C6, 99% (Cambridge isotope laboratories, CLM-2265-H-0.1) and 40mg/ml L-lysine:2HCL 4,4,5,5-D4, 96-98% (Cambridge isotope laboratories, DLM-2640). After co-cuture, T cells were washed away, spun down at 1700 rpm for 5 min and washed with PBS. Tumor cells were washed 2x PBS, scrapped in 1 ml PBS and pelleted by spin down at 1500 rpm for 5 min and combined together with the corresponding T cell samples recovered from the media.

### HySic mass spectrometry sample preparation

Cell pellets were resuspended in 300 uL of 1%(w/v) sodium deoxycholate (SDC) lysis buffer (100mM Tris pH 8.0, 10mM Tris (2-carboxyethyl) phosphine (TCEP), 40mM chloroacetamide, phosphatase inhibitor (PhosSTOP, Roche), and protease inhibitor (complete mini EDTA-free, Roche). Resuspended samples were boiled for 5 min at 95°C and sonicated for 15 min in the Bioruptor (Diagenode) at 30 sec on, 30 sec off cycles. Lysates were clarified by centrifugation (20,000 x g for 10 min) and protein concentration was determined using a Bradford protein assay. For each sample, 210 ug of protein was digested using Lys C (Wako) at an enzyme:protein ratio of 1:100 and trypsin (Sigma) at 1:50 ratio overnight at 37 °C. Samples were acidified by addition of formic acid (FA) to a final concentration of 2% (v/v). Samples were centrifuged (20,000 x g for 10 min) and desalted using 5 uL C18 cartridges on the AssayMap BRAVO platform (Agilent Technologies). For each sample, 10 ug of digested protein was saved for proteome analysis and the remaining 200 ug of digested protein was enriched for phosphopeptides using 5 uL Fe(III)NTA cartridges on the AssayMap BRAVO platform, as described previously (Post et al., 2017). Briefly, samples were dissolved in 200 µL loading buffer (80% acetonitrile (ACN) / 0.1% TFA). Fe(III)-NTA 5 µL cartridges were primed with 200 µL of 0.1% TFA in ACN and equilibrated with 250 µL of loading buffer. Then, the column was loaded the samples at a loading speed of 5 µL/min. The columns were washed with 250 µL loading buffer and eluted with 35 µL of 5% ammonia solution into 35 µL of 10% FA. All elusions were dried in a vacuum centrifuge and resuspended in 2% FA for LC-MS injection. For phosphopeptide elutions, samples were injected twice when possible.

### Generation of ‘TO’ control samples

For ‘T0’ samples, ^8^K^10^R labeled tumor cells were incubated for four hours in ^4^K^6^R media, without T cells, to establish an internal control for background translation. Following cell lysis, tumor T0 samples were mixed with lysed T cells (labeled ^0^K^0^R) to create an artificially mixed proteome sample. This mixed T0 sample possessed a comparable MS1 profile to 2h, 4h, and 6h co-culture lysates, which was necessary for subsequent LFQ analysis.

### HySic mass spectrometry data acquisition

Proteome and phosphoproteome data was acquired on either an Orbitrap Exploris 480 MS (Thermo Scientific) coupled to an UltiMate 3000 UHPLC (Thermo Scientific) fitted with a C18 trapping column (PepMap100, 5um, 100A, 5mm x 300um; Thermo Scientiifc) and a homemade C18 analytical column (120 EC-C18, 2.7uM, 50 cm x 75um; Agilent Poroshell) or acquired on an Orbitrap Q Exactive H-FX MS (Thermo Scientific) coupled to an Agilent 1290 Infinity UHPLC system fitted with a Reprosil pur C18 trap column (100um x 2cm, 3um, Dr. Maisch) and a homemade C18 analytical column, described above. The different instrumentation used corresponds to the two separate experimental batches (HySic Study I: A549 and NCI-H358 cell lines, H-FX-Agilent 1290 setup; HySic study II: LCLC-103H and EBC-1 cell lines, Explores 480-UltiMate setup). The LC-MS parameters for each experiment were as follows: SI proteome data was collected over a 175 min run following 5 min of trapping with buffer A (0.1% FA). Peptides were eluted over a 155 min gradient from 10% to 44% solvent B (0.1%FA, 80%ACN) at a flow rate of 300 nL/min, followed by a wash and re-equilibration step. MS data was acquired in Data-Dependent mode with MS1 settings at 60,000 resolution, scan-range 375-1600 m/z, AGC target at 3E6 and max ion injection time (IT) 20 msec. HCD-MS2 spectra were collected at 30,000 resolution for the top 15 precursors using a NCE of 27, max IT of 65 msec, AGC target of 1E5, isolation window of 1.4 Da, fixed first mass set at 120 m/z, and a dynamic exclusion time of 30 sec. Study I phosphoproteome data was collected for 115 min using a 9%-36% solvent B elution gradient over 95min. MS settings were identical as above with the following adjustments: the top 12 precursors were selected for fragmentation with a max IT of 85 msec and dynamic exclusion of 18 sec. Study II proteome and phosphoproteome data was collected using settings identical to Study I with the following exceptions: AGC targets were set to ‘standard’ and max IT was set to ‘auto’. MS2 scans were collected for a fixed cycle window of 3 sec with an NCE of 28 and a dynamic exclusion time of 24 sec for proteome and 16 sec for phosphoproteome data.

### HySic mass spectrometry data processing

MS raw files were searched using MaxQuant software (versions 2.0.1.0 and 1.6.10.43) against a reviewed homo sapiens database (UniProt, 2017; 21,008 entries) and a contaminant database. Study I (A549 and NCI-H358 tumor cell lines) and Study II (LCLC-103H and EBC-1 tumor cell lines) raw files were processed separately and in each case proteome and phosphoproteome data was searched separately, generating a total of 4 datasets. For each MaxQuant search, labeling multiplicity was set to 3 with Arg6, Lys4 and Arg10 and Lys8 selected for medium and heavy labels, respectively. Match-between-runs was enabled, Re-quantify was disabled, fixed modifications were set to carbamidomethylation, variable modifications were set to methionine oxidation and protein N-term acetylation, and all other MaxQuant parameters were set to default. For phosphoproteome searches, phosphorylation at STY residues was selected as an additional variable modification. Protein and phosphosite intensities were extracted from MaxQuant output files for each SILAC label independently (’H’, ‘M’, ‘L’). All SILAC ratio data was discarded. Extracted protein and peptide intensities were further processed in R using in-house scripts. Within each experimental dataset, the total summed intensity for each raw file were compared to identify sample outliers (i.e. failed sample injections). An outlier was defined as a sample with a signal intensity > 2x standard deviation below the mean of all samples. No outliers were identified in either proteome dataset, however, several injections were discarded from phosphoproteome data (**Fig. S. 1C**). Next, to correct for differences in sample loading or instrument performance, samples were median normalized based on the combined intensities of all SILAC channels in each sample. Injection replicates were averaged to generate one dataset per biological replicate. After this step, SILAC intensities for each sample were processed independently, generating a total of 3 data tables for each experimental dataset: tumor cell, T cell, and Newly Synthesized proteins. Extracted T cell and tumor cell data was median normalized again to correct for any differences in cell mixing across co-culture samples (**Fig. S. 1D**). Newly Synthesized proteins did not undergo a second normalization because these abundances are not expected to be similar across all samples, but rather increase with time (**Fig. S. 1D**).

### HySic mass spectrometry data filtering

All datasets were filtered to require at least 2 biological replicates (BR) in at least one experimental condition. Further filtration steps were applied per dataset as described below. Phosphoproteome data for tumor cells and T cells was filtered to required phosphosite quantification in at least 2/3 BRs for one or more co-culture incubation time points (2h, 4h, 6h). For heat map analysis, phosphosite changes relative to time point zero (T0) were calculated to identify phosphorylation changed induced by co-culture conditions. Within each tumor cell line co-culture series, T0 values were averaged and phosphosite abundances were normalized to T0 averages. For T cell calculations, T0 values were averaged within experimental batches, as these T cells were aliquots from the same donor. For phosphosites with no T0 values in any BR, data was imputed using the minimum value from the dataset. T0 ratio data was median normalized and an ANOVA test was performed to identify significantly changing phosphosites within each coculture treatment series (P value < 0.05). Phosphosites with no T0 values were also included in the ‘significant hits’ list as they may represent ‘on/off’ phosphorylation states. Finally, data was filtered for phosphosites that show significant changes within at least three of the four co-cultured cell lines.

Proteome data for tumor and T cells was filtered to require measurements at T0 and 2h time points. For heat map analysis, T0 ratios were calculated as described above, and data was filtered to require at least 2/3 BR at the 2h timepoint and significant downregulation relative to T0 (p < 0.05) in at least one cell line. Data was further filtered to include proteins that were decreasing in at least three of the four co-cultured cell lines at the 4h and 6h timepoints. For newly synthesized proteins, proteome data was filtered to require quantification in at least 2/3 BR in the 4h and/or 6h co-culture time points. Proteins were further filtered to require either no measurement in T0 controls for background translation or have at least a 2-fold increase above background translation at 4h and 4-fold increase above background translation at 6hr. For newly synthesized phosphoproteome data, similar filtering steps were applied. For inclusion in heat map analysis, phosphosites must also show quantification at 6h in at least three of the four co-cultured cell lines.

### T cell PAK1 inhibitor MS sample preparation

T cell pellets were lysed in a 1% SDC buffer, quantified, digested (60 ug/sample), desalted, and dried as described above. Digested samples were resuspended in 80 uL of 50 mM HEPES buffer and labeled with 10-plex TMT reagents (Thermo Fisher Scientific, LOT# UA275089) at a label:protein ratio of 2:1 for 1.5 h, shaking at room temperature. To check labeling efficiency and mixing ratios, 2 µl of each sample was combined in acidic solution for an LC-MS test run. After complete labeling was confirmed, samples were quenched using 6 uL of 5% hydroxylamine and incubate 15 min at room temperature. Labeled samples were next mixed equally and desalted using SepPak C18 cartridges (Waters). Alongside the individual samples, a pooled reference sample was generated from an aliquot of all digests. This pooled reference sample labeled was spiked into each TMT-10plex pool to act as a universal standard for data normalization. Following desalting, an aliquot of each TMT pool was reserved for proteome analysis and the rest was dried in the vacuum-centrifuge for phosphopeptide preparation. The workflow followed for the TMT sample preparation is outlined in (**Fig. S. 7A)**. Before enrichment of phosphopeptides, the TMT labeled samples were fractionated using reversed-phase S cartridges and high pH buffer (200mM NH_4_HCO_3_, pH 10) on the AssayMap BRAVO Platform. Fractions were eluted with four steps of increasing buffer B (100% ACN): 22%, 30%, 36%, and 70% directly into 10% FA to neutralize the pH. Next, phosphopeptides were enriched for each fraction on the AssayMap BRAVO platform following the method described above.

### T cell PAK1 inhibitor MS data acquisition

TMT labeled samples for proteome analysis were measured using an Orbitrap Fusion Lumos Tribrid MS (Thermo Scientific) coupled to an UltiMate 3000 UHPLC system (Thermo Scientific) fitted with a µ-precolumn (C18 PepMap100, 5 µm, 100 Å, 5 mm × 300 µm; Thermo Scientific), and a homemade analytical column (120 EC-C18, 2.7 µm, 50 cm × 75 µm; Agilent Poroshell). Samples were resuspended in 2% FA solution and loaded in solvent A (0.1% FA in water) with a flow rate of 30 µl/min and eluted using a 175 min gradient at a flow rate of 300 nL/min. The gradient for peptides was as follows: 9% solvent B (0.1% FA in 80% ACN) for 1 min, 9-13% B for 1 min, 13-40% B for 155 min, followed by a was hand re-equilibration step. The MS was operated in TMT-SPS-MS3 mode. The following MS settings were applied. For MS1 scans, detector type: Orbitrap; resolution: 60,000; scan range (m/z): 375-1500 Th; AGC target: standard; maximum injection time mode: auto; intensity threshold: 5 x10^3^; charge state: 2-6; dynamic exclusion: 30 s. For MS2 scan, isolation mode: quadrupole; isolation windows (m/z): 1.2 Th; CID collision energy: 35%; precursor selection mass range (m/z): 400-1600 Th; precursor ion exclusion mass width: low 25 ppm, high 25 ppm; isobaric tag loss exclusion: TMT. For SPS-MS3 scan, number of SPS precursors: 5; MS isolation window (m/z): 1.3 Th; activation type: HCD; collision energy: 65%; detector type: Orbitrap; resolution: 50,000; scan range: 100-500 Th; normalized AGC target: 200%; maximum injection time: 200 ms.

TMT labeled phosphopeptide samples were measured using MS2-based quantification on an Orbitrap Exploris 480 mass spectrometer (Thermo Fisher Scientific) coupled to an UltiMate 3000 UHPLC system (Thermo Fisher Scientific) as described above. Data was acquired using a 115 min acquisition method, as described previously for the HySic phosphoproteome, with the following exceptions: MS2 resolution was set to 45,000, MS2 isolation window was reduced to 1.2, and NCE was increased to 32.

### T cell PAK1 inhibitor MS data processing

Raw data files were processed with Proteome Discover 2.4 (Thermo Fisher Scientific) using Sequest HT and with a Swiss-Prot Homo sapiens database (version. 20220720) and a contaminant database. The isotopic impurities of the TMT reagent were corrected using the values specified by the manufacturer. The search parameters for proteomic data were as follows: enzyme was set to trypsin, with up to 2 missed cleavage sites. Fragment mass tolerance was set to 0.5 Da and precursor mass tolerance to 10 ppm. Carbamidomethyl of cysteine and TMT10-plex modification of lysine and peptides N-terminus were set as static modifications. Oxidation of methionine and protein N-terminal acetylation were set as dynamic modifications. The reporter ions quantifier was set with HCD and MS3 (mass tolerance, 20 ppm). The PSM validation was performed using Percolator and the false discovery rate was set to 0.01. For the reporter ion quantification, unique + razor peptides were used. The threshold of co-isolation, average reporter S/N and SPS mass matches were 45, 10 and 65% respectively. The site probability threshold of peptide group modification was set 75. The FDR of peptide validator for PSMs and peptides were set to 1%. The FDR for protein FDR validator was also set to 1%. Minimum peptide length was set to 6 and minimum number of peptide sequence was set to 1 for peptide and protein filter. The parameters for phosphopeptide searches were as follows: Enzyme was set to trypsin, with up to 2 missed cleavage sites. Fragment mass tolerance was set to 0.06 Da and precursor mass tolerance to 20 ppm. Carbamidomethyl of cysteine and TMT10-plex modification of lysine and peptide N-terminus were set as static modifications. Oxidation of methionine and protein N-terminal acetylation were set as dynamic modifications. Phosphorylation of serine, threonine and tyrosine was set as dynamic modifications. The reporter ions quantifier was set with HCD and MS2 (mass tolerance, 20 ppm). The PSM validation was performed with Percolator and the false discovery rate was set to 1%. For the reporter ion quantifier, unique + razor peptides were set for the quantification. The threshold of co-isolation, average reporter S/N and SPS mass matches were 45, 10 and 65% respectively. The site probability threshold of peptide group modification was set 75. The FDR of peptide validator for PSMs and peptides were set to 0.01. The FDR for protein FDR validator was also set to 0.01. Minimum peptide length was set to 6 and minimum number of peptide sequence was set to 1 for peptide and protein filter.

### Flow cytometry

For flow cytometry analysis of T cell:tumor co-cultures, 5×10^4^ tumor cells were co-cultured with 5×10^4^ T cells (1:1 ratio) in a round bottom 96 well plate (Greiner, M9436-100EA) for 6 hours. For intracellular staining, GolgiPlug^TM^ Protein Transport Inhibitor (1:1000, BD Biosciences, BD555029) was added after 3h co-culture. The Foxp3 Transcription Factor Staining Buffer kit (Invitrogen, 00-5523-00) was used following manufacturer instructions. For cell surface staining, cells were washed and stained with antibodies in 0.1% BSA PBS buffer on ice for 30 min protected from light. After staining, cells were washed twice and analyzed using LSRFortessa (BD Biosciences). Cells were gated on FSC and SSC followed by single cell gate on FSC-HH/FSC and SSC-H/SSC and a LIVE/DEAD near IR marker (1:1000, BD Biosciences, L34976). The following antibodies were used: CXCL-9-APC (1:20, Biolegend, 357906), PD-L1/CD274-PECy7 (1:50, BD Bioscience, 558017), IFN-y-APC (1:100, BD, 554702).

For flow analysis of T cells treated with NVS-PAK1-1 or DMSO, cell surface or intracellular staining was performed as described above and the following antibodies were used: LIVE/DEAD Yellow marker (1:1000, Themo Fisher, L3496), PD1-BV650 (1:100, Biolegend, 329739), CD137-APC (1:100, BD Biosciences, 550890), CD69-PE (1:100, Immunotools, 21620694), CD25-PerCPCy5.5 (1:100, Biolegend, 302625) TNFa-PE (1:100, Biolegend, 502909), IFN-y-APC (1:100, BD Biosciences, 554702) and CD8-Pacific Blue (1:100, Biolegend, 344717).

### Western Blotting

For western blot analysis of T cell:tumor co-cultures, cells were seeded in a 10 cm petri dish. After co-culture T cells were collected from the supernatant and spun down at 1700 rpm for 5 min. Tumor cells were washed twice with PBS and spun down at 1500 rpm for 5 min. Cell pellets were then resuspended in RIPA lysis buffer supplemented with protease/phosphatase inhibitor cocktail (1:1000, Thermo Fisher, 78440) and lysed for 30 min on ice. Protein concentration was quantified using Bio-Rad protein assay (Bio-Rad, 500-0006). Samples were analyzed on NuPAGE Bis-Tris 4-12% polyacrylamide-SDS gels (Life technologies, NP0321BOX) in MES buffer (Invitrogen, B000202). They were subsequently transferred into nitrocellulose membranes using iBlot^TM^ (Thermo Fisher). Membranes were blocked for 1 hour with 4% milk in PBST (0.2% Tween-20 in PBS). Primary antibodies were incubated overnight at 4°C. The next day membranes were washed with PBST and incubated with the secondary antibodies for one hour at RT. After final wash in PBST, membranes were developed with Super Signal West Dura Extended Duration Substrate (Thermo Fisher, 34075) in ChemiDoc (Bio-Rad). The following antibodies were used: IFNyR1 (1:1000, Santa Cruz, sc-28363), totalSTAT1 (1:1000, Cell Signaling, 14994), p-STAT1 Ser727 (1:1000, Cell Signalling, 9167), Vinculin (1:1000, Cell Signaling, 4650), Tubulin (1:1000, Sigma Aldrich, T9026), p-PAK1 S144 (1:1000, CST, 2606T,), totalPAK1 (1:500, Biolegend, 868601), peroxidase conjugated anti-mouse IgG (1:5000, Thermo Fisher, G-21040) and anti-rabbit IgG (1:5000, Thermo Fisher, G-21234).

### PAK1 inhibitor treatment

For NVS-PAK1-1 (MedChem, HY-100519) treatment in co-culture experiments, 1-4×10^4^ tumor cells were seeded in flat bottom 96 well plates (Greiner, 655180) with the indicated ratios of T cells. NVS-PAK1-1 was added at the indicated concentrations. Cell viability was determined using the Incucyte Zoom (HERACELL 240i incubator, Essen Bioscience) every 2 hours by counting red fluorescence cells (mPlum expression of tumor cells). Cell count was normalized by subtraction of the cells counted at time 0h.

For NVS-PAK1-1 treatment of T cells, cells were re-activated in a non-culture treated 24 well plate that was coated with aCD3 and aCD28 (1:200 in PBS, Invitrogen, 16-0037-85) and aCD28 (1:200 in PBS, Invitrogen, 16-0289-85) for 24 hours. Cells were then stained with CFSE (1:6000 in PBS, Biolegend, 423801) for 20 min at 37°C and washed with 2% BSA in FBS. T cells were then maintained at 10^6^ cells/ml with NVS-PAK1-1 or DMSO for the indicated times. Cell surface markers were analyzed by flow cytometry as described above.

### Bioinformatic analysis

For pathway enrichment analysis g:Profiler (Raudvere et al., 2019) and Metascape (Zhou et al., 2019) were used. Reactome, Wikipathways, KEGG, Hallmark and GO significant (p<0.05) pathways were shown when indicated.

### Statistical analysis

All experimental data was tested for normal distribution with Saphiro-Wilk test. Datasets that passed normality were analyzed with parametric tests and datasets that did not pass were analyzed with non-parametric. In each figure legend the statistical test is indicated. Data were analyzed and plotted in Graphpad Prism. Proteome and phosphoproteome data were analyzed using R and statistical significance was determined used ANOVA analysis.

## Acknowledgements

We would like to thank the Schumacher lab for sharing the MART-1 TCR system. We thank our colleagues in the lab and Division for valuable input and discussion and Nils Visser for technical support. We would also like to thank the flow cytometry facility in NKI-AVL for contributing to this work. This work received support from the Horizon 2020 program INFRAIA project Epic-XS (Project 823839) and the NWO funded Netherlands Proteomics Centre through the National Road Map for Large-scale Infrastructures program X-Omics (Project 184.034.019). DSP is funded by, and a member of, the Oncode Institute, which is partly financed by the Dutch Cancer Society.

## Author contributions

S.I.M., M.A., D.S.P. and K.S. conceived the study and designed the experiments. S.I.M., J.P. and A.A. performed cell culture experiments. F.W. and K.S. performed mass spectrometry analysis. S.I.M. and K.S. performed bioinformatics analysis. S.I.M., D.S.P. and K.S. wrote the manuscript. All authors read and approved the manuscript. The project was supervised by M.A. and D.S.P.

## Data availability statement

Data available within the article or its supplementary materials. All mass spectrometry proteomics data have been deposited to the ProteomeXchange Consortium via the PRIDE (Perez-Riverol et al., 2022) partner repository with the dataset identifier PXD041526.

---

**S.1 Supplementary to figure 1.**
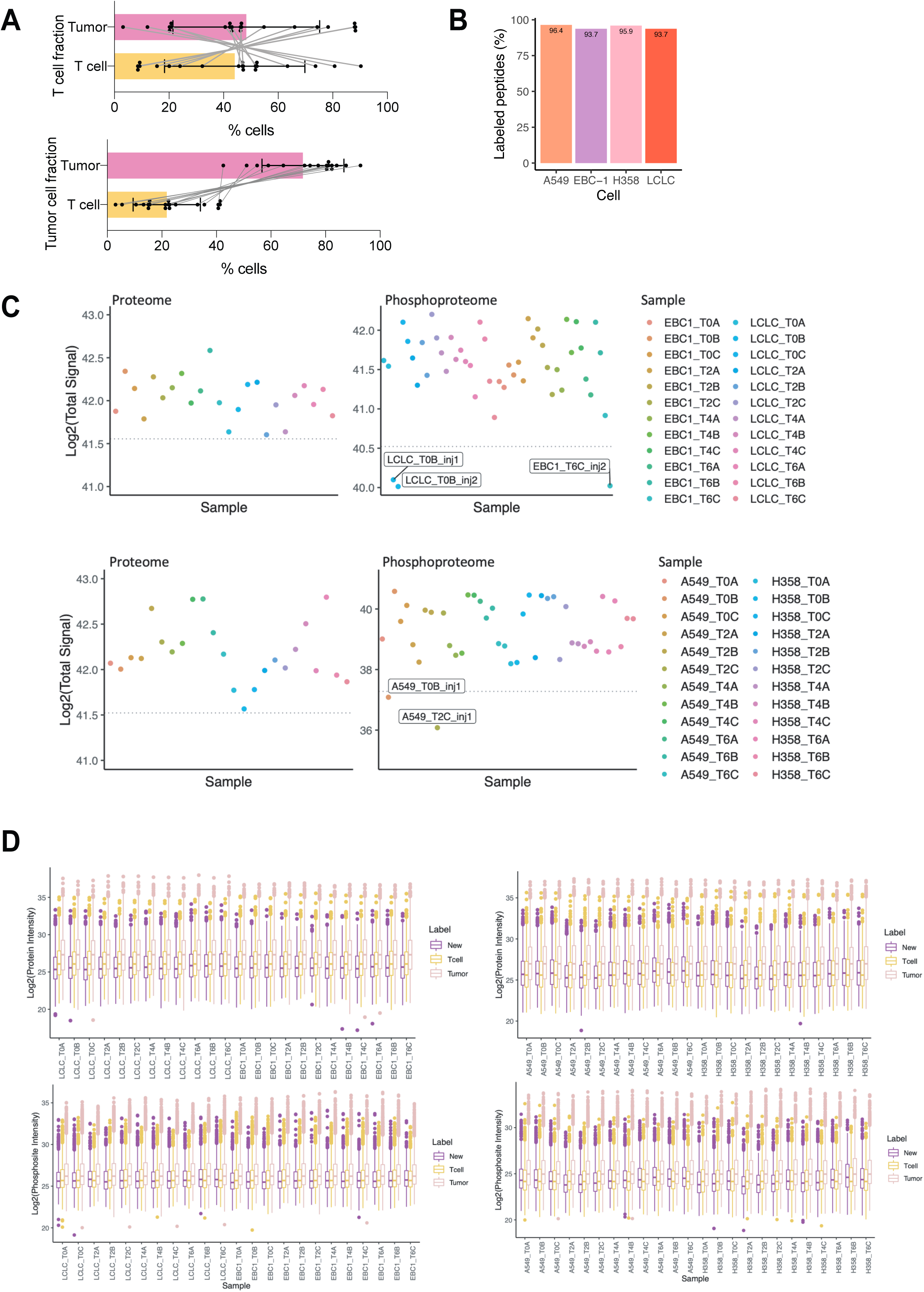
A) Percentage of cells measured by flow cytometry after co-culture in T cell (top) or tumor cell (bottom) fractions. Tumor cell percentage is represented in pink bar, T cell percentage is represented in yellow bar. Each dot represents a cell line. Grey lines connect the percentage of each cell types in individual cell lines. B) SILAC labeling efficiency check. Bars represent heavy labeled peptide incorporation in each tumor cell line. Tumor cell lysates were analyzed by LC-MS and data was searched with K8 and R10 as a variable modification. Labeled and unlabeled peptides were counted. LCLC is abbreviation of LCLC-103H, H358 is abbreviation of NCI-H358. C) Sample outlier detection. Y axis represents log2 summed abundance for each sample. Dotted line represents 2x standard deviation from the mean of the entire dataset. Samples below the line were removed from the study. Phosphoproteome samples had two replicate injections on the MS. LCLC is abbreviation of LCLC-103H, H358 is abbreviation of NCI-H358. D) Normalized protein and phosphosite expression levels. Y axis represents median log2 abundance for each sample. Tumor and T cell samples were normalized across all samples in the batch. Phosphoproteome injection replicates are averaged into generate one sample per biological replicate. LCLC is abbreviation of LCLC-103H, H358 is abbreviation of NCI-H358.

**S. 2 Supplementary to figure 2.**
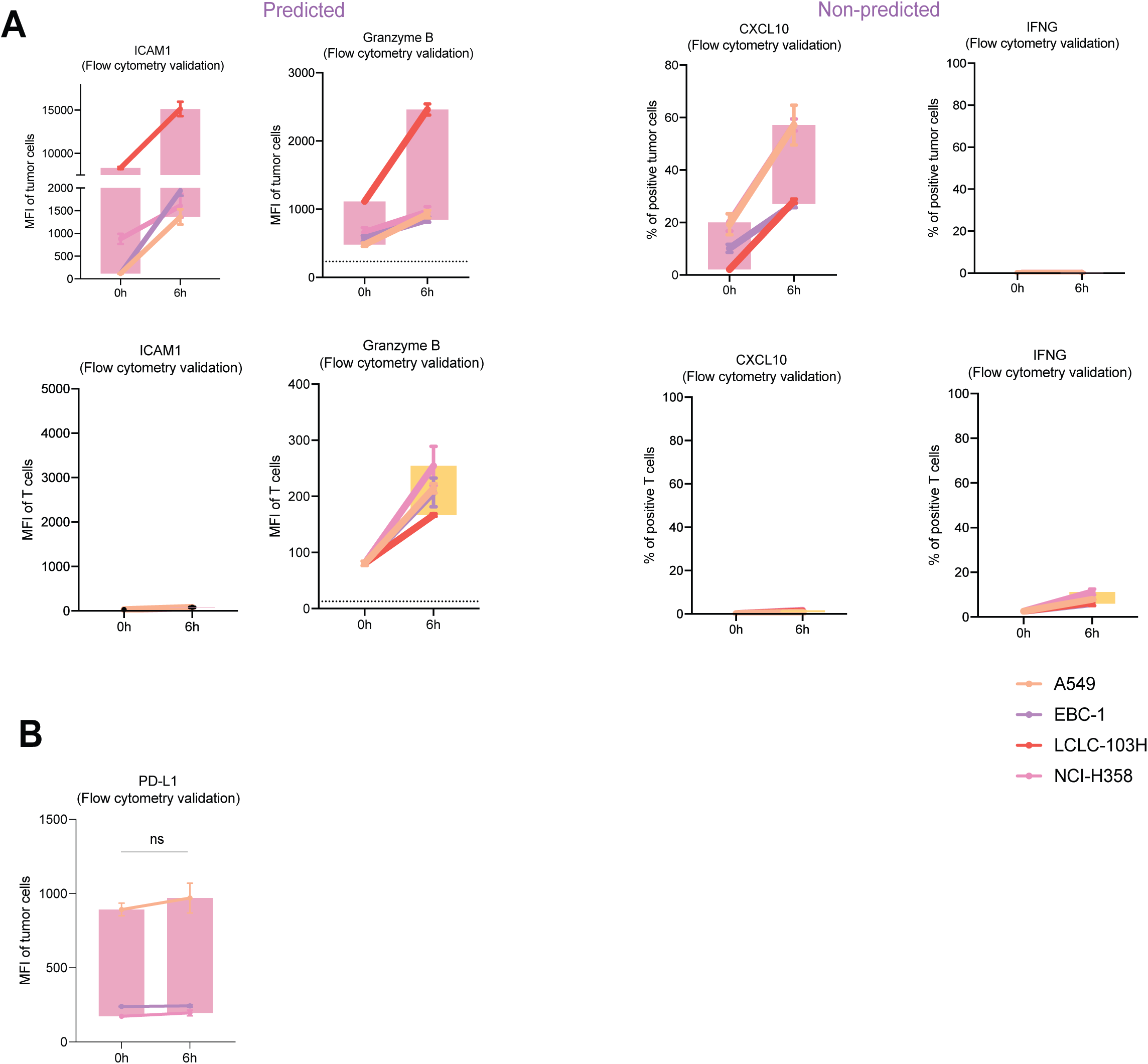
A) Independent T cell donor experiment of Fig. 2E. Flow cytometry protein expression of the indicated proteins in either tumor cells (pink boxplots) or T cells (yellow boxplots). Each cell line is represented by a colored dot with SD bars. Boxplot indicates deviation of four cell lines combined. **p-value<0.01, ****p-value<0.0001. 2-way ANOVA test used for statistical analysis. # indicates measurement below detection levels. B) Flow cytometry expression of PD-L1 protein in tumor cells across three independent cell lines. ns indicates not significant.

**S. 3 Supplementary to figure 3.**
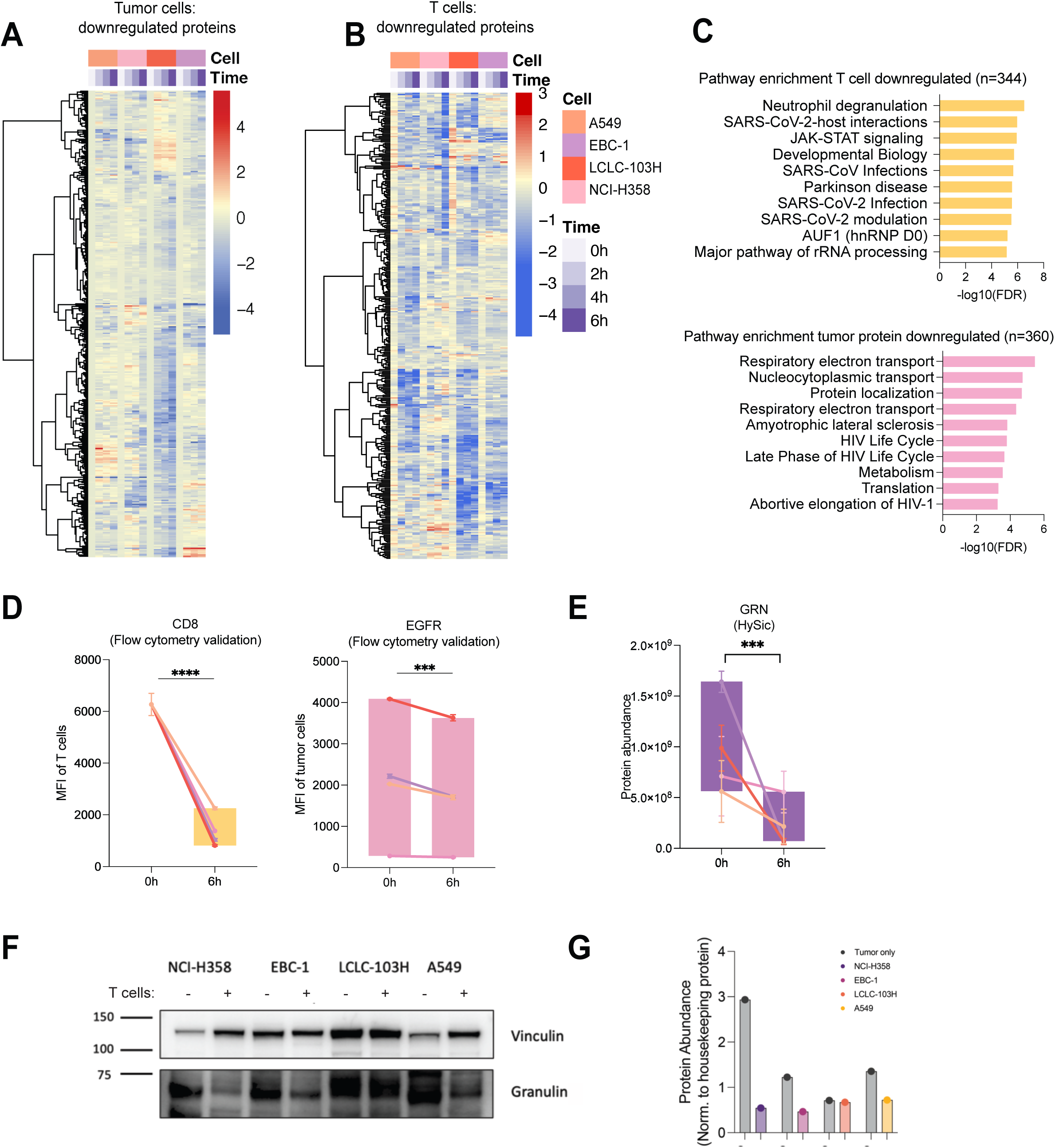
A) Heatmap of significantly downregulated tumor cell proteins (p< 0.05) relative to T0 at any co-culture timepoint in at least one co-cultured cell line. Proteins included in the heatmap were required to show a decreasing trend in at least 3/4 cell lines. B) Heatmap of significantly downregulated T cells proteins (p< 0.05) relative to T0 in at any co-culture timepoint in at least one co-cultured cell line. Proteins included in the heatmap were required to show a decreasing trend in at least 3/4 cell line co-cultures. C) Top 10 enriched pathway in downregulated proteins detected in T cell (top) or tumor (bottom) channel. Enrichment was performed using *Gprofiler* with Reactome and WikiPathways databases. Downregulated proteins were selected from heatmap data in A & B. Pathways are listed from smallest to largest *p*-value. D) Independent T cell donor experiment of Fig. 3C. Median fluorescence intensity (MFI) measured by flow cytometry of the indicated proteins in either tumor cells (pink boxplots) or T cells (yellow boxplots). Each cell line is represented by a colored dot with SD bars. Boxplot indicates deviation of four cell lines combined. ***p-value<0.001, ****p-value<0.0001. 2-way ANOVA test used for statistical analysis. E) Protein abundance detected by HySic of GRN. ***p-value<0.001. Mixed model analysis performed because of absence of one data point (non-detected in MS). F) Measurement of Granulin protein expression by western blotting in the indicated tumor cell lines after 4h of co-culture with or without (-) T cells. G) Quantification of F normalized to vinculin.

**S. 4 Supplementary to figure 4.**
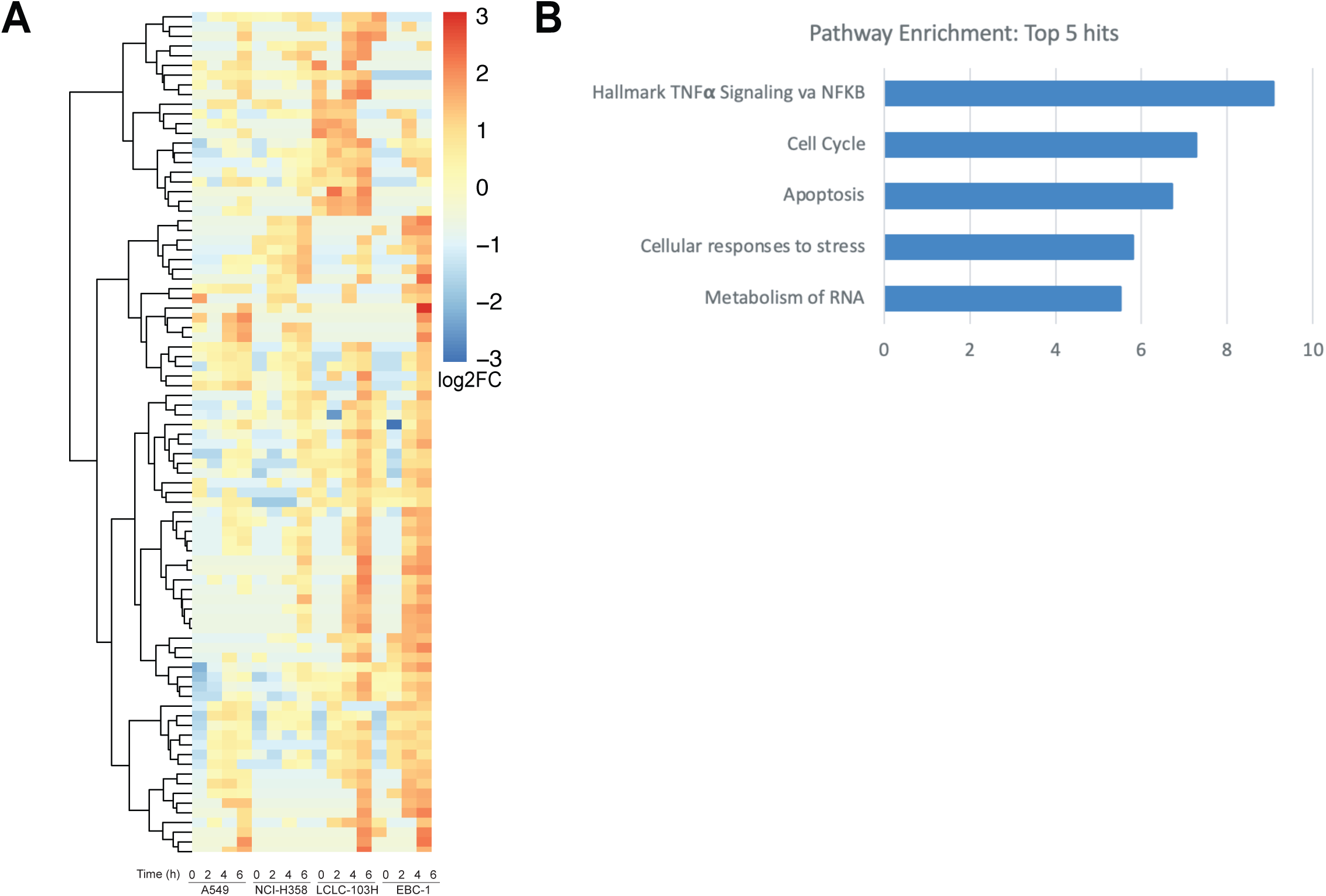
A) Heat map of upregulated phosphosites in newly synthesized proteins in at least 3/4 co-cultured cell lines. T0 normalized data wherein samples with no T0 detection were imputed with the minimum value from the dataset. Missing values were replaced with zero for heatmap visualization. B) Pathway enrichment (Metascape) for upregulated phosphorylated proteins from newly synthesized phosphosite heatmap. Top five enrichment terms are displayed.

**S. 5 Supplementary to figure 4.**
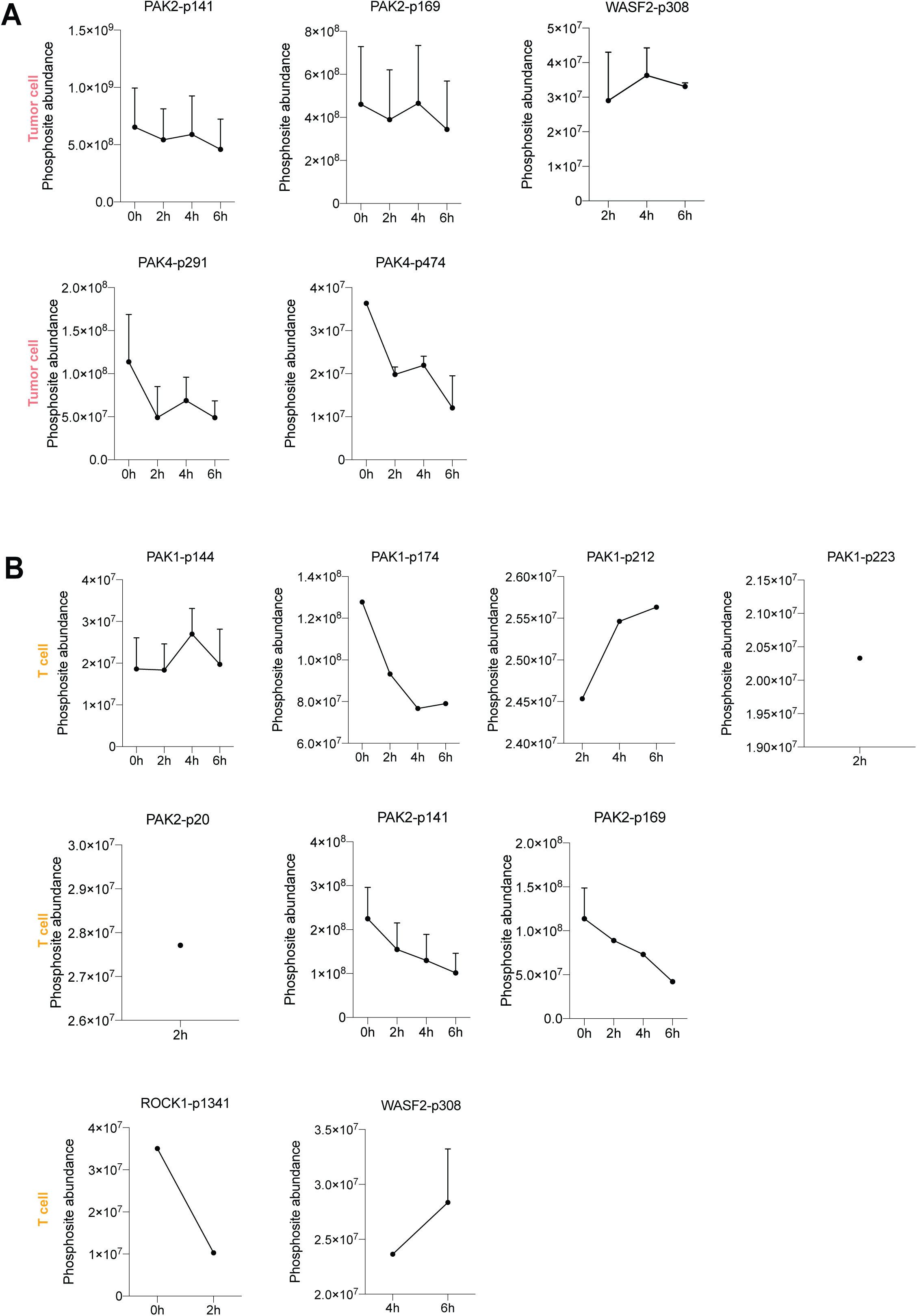
A) Abundance of the indicated phosphorylation site in tumor cells detected by HySic. Each dot represents average of four cell lines and error bars represent SEM. B) Abundance of the indicated phosphorylation site in T cells detected by HySic. Each dot represents average of four cell lines and error bars represent SEM.

**S. 6 Supplementary to figure 4.**
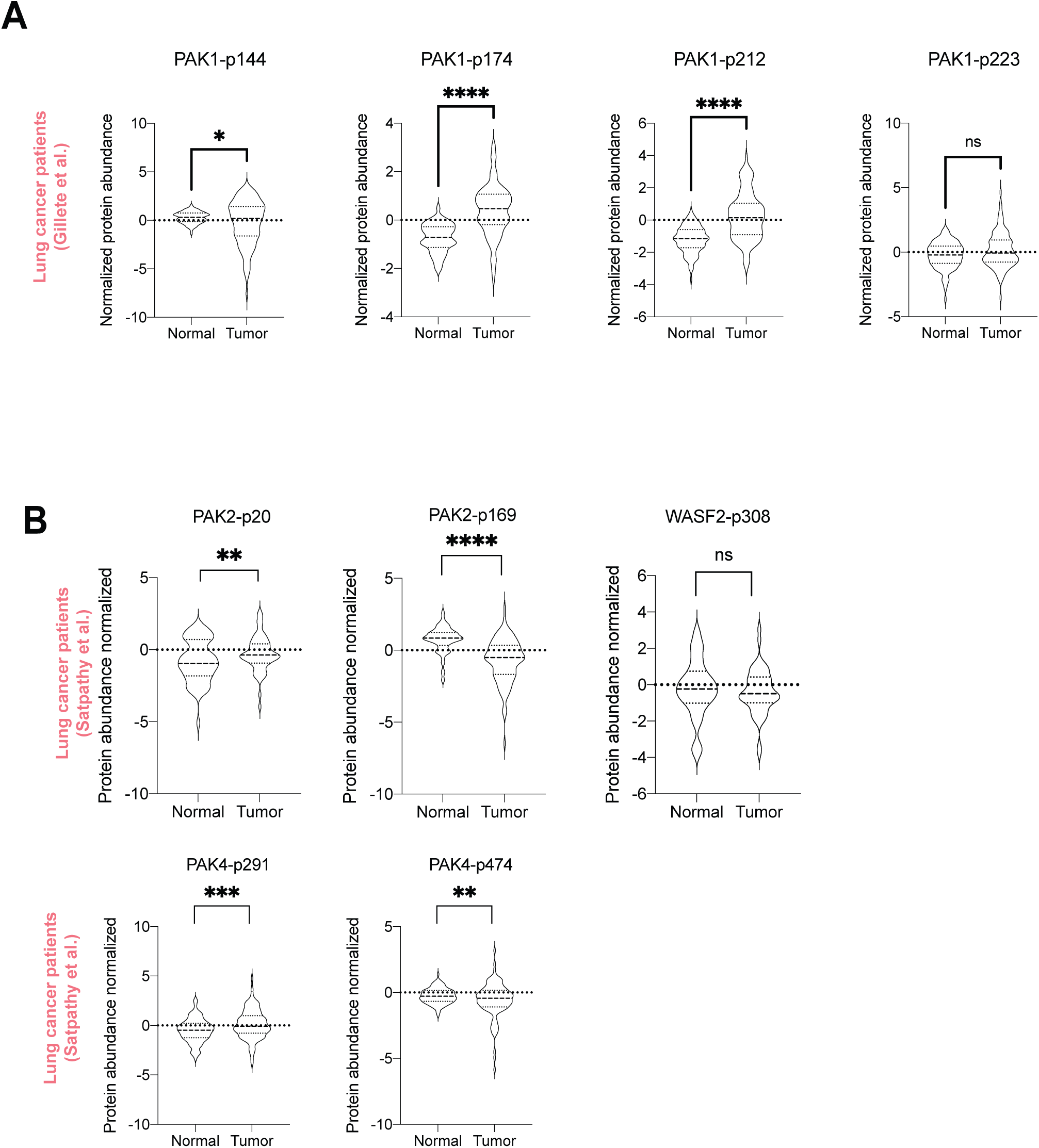
A) Normalized phosphosite abundance from Satpathy *et a*l. of the indicated proteins in normal versus tumor tissue. Violin plots represent average of individual patient expression levels. B) Normalized phosphosites abundance from Gillete *et al*. of the indicated protein sites in normal versus tumor tissue. Violin plots represent average of individual patient expression levels.

**S. 7 Supplementary to figure 5.**
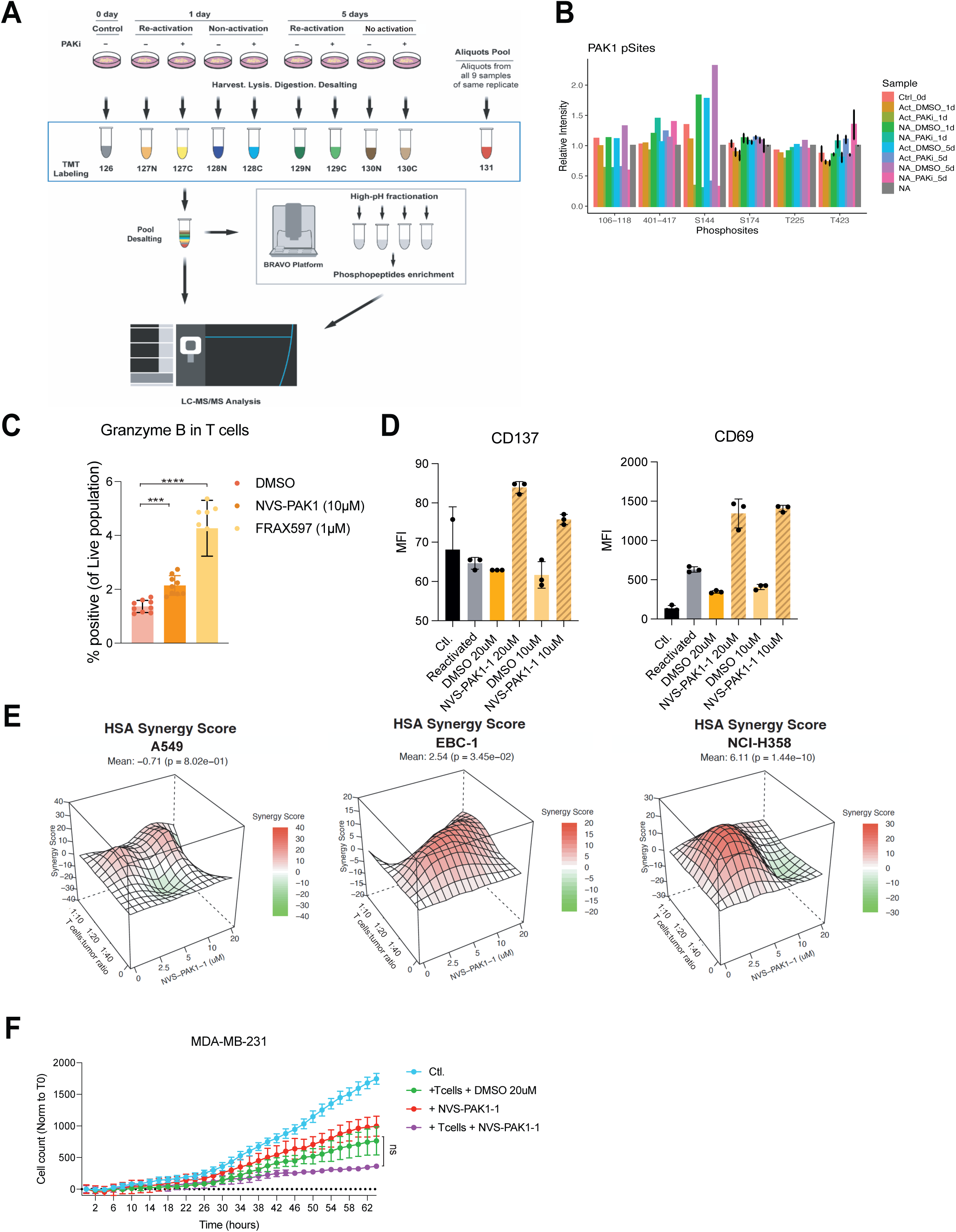
A) Experimental workflow for TMT labeling and sample handling for PAK1 inhibitor treated T cells. B) Phosphoproteomic analysis of T cells treated with DMSO or NVS-PAK1(10uM) for 1 or 5 days. T cells were activated (ACT) or not (NA). The relative intensities of the indicated PAK1 phosphosites are shown. Error bars represent standard deviation from 3 biological replicates. C) Independent T cell donor experiment of Fig. 5B. Percentage of Granzyme B-positive T cells (of live population) upon NVS-PAK1 (10uM), FRAX587 (1uM) or DMSO treatment for 5 days. ***p-value<0.001, ****p-value<0.0001. 2-way ANOVA test used for statistical analysis. D) Measure of protein expression by median fluorescence intensity (MFI) of CD137 or CD69 surface markers in flow cytometry of T cells treated with NVS-PAK1 (10uM or 20uM) or DMSO for 5 days. E) Synergy score (HSA) calculated with *Synergyfinder* on the indicated T cell:tumor cell ratios and drug concentrations for A549, EBC-1 and NCI-H358 cell lines. Score indicates the percentage of additive effect of using a combination of two treatments compare to the single agent. F) Cytotoxic assay of tumor cells and T cells co-cultured with or without NVS-PAK1 in Incucyte®. Inhibitor concentration and T cell:tumor cell ratio was optimized per cell line; MDA-231 (10uM, 1:40). Data was normalized to 0h. Error bars represent SD of 3 technical replicates. Statistical analysis was performed by Friedman test. ns indicate not significant.

**S. 8 Supplementary.**
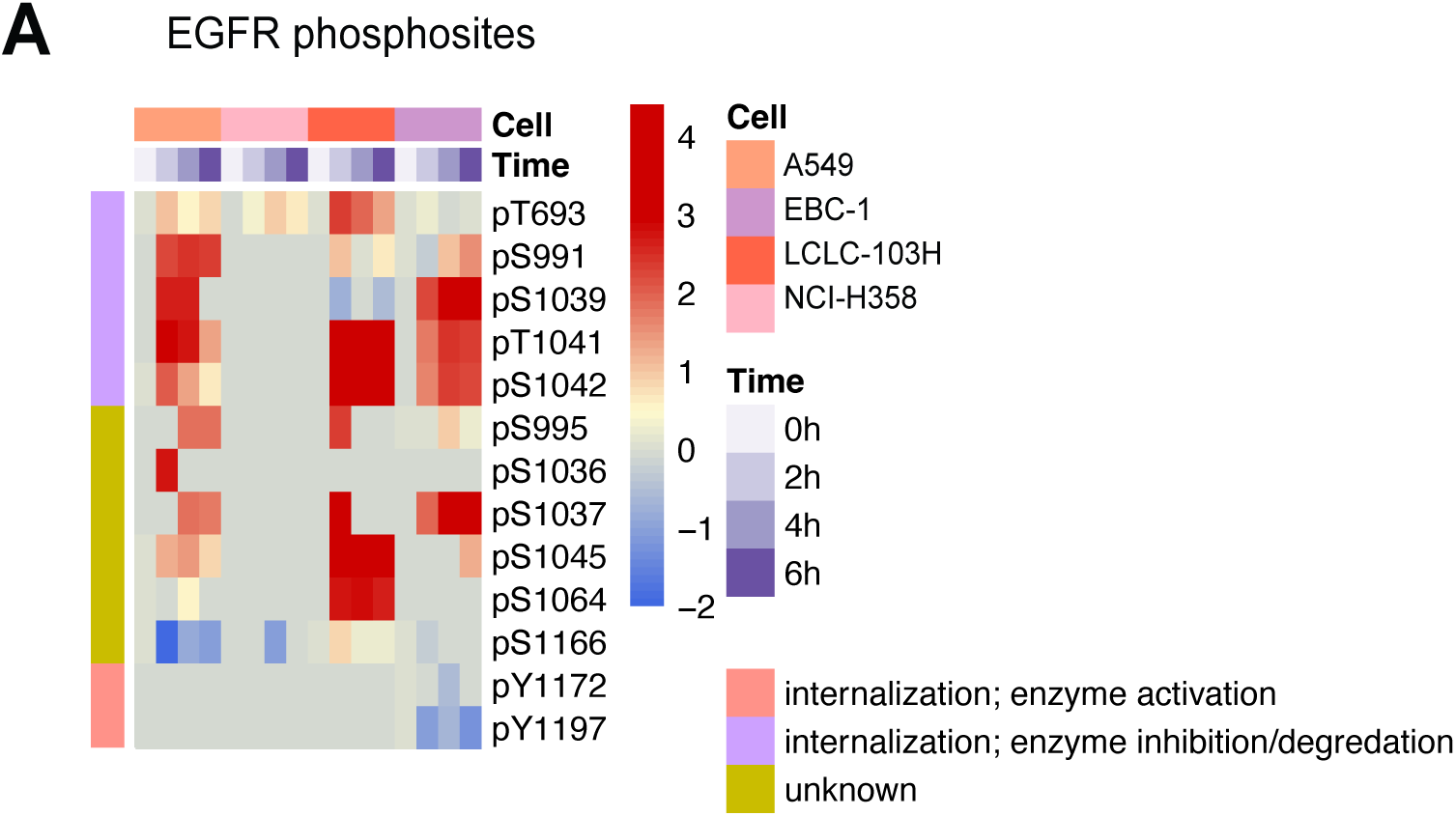
A) Heatmap clustering of EGFR phosphorylation sites. Phosphosite abundances are normalized to T0 values and log2 transformed. Missing values are replaced with zero for heatmap clustering. Phosphosite functional annotations were extracted from PhosphoSitePlus.

